# Structural basis of underwound DNA topology bearing PAM-mutant recognition by AtCas9

**DOI:** 10.64898/2026.01.30.702744

**Authors:** Min Duan, Bing Meng, Lei Zhou, Lijie Wu, Xiaohan Tong, Dongchao Huang, Hao Yin, Zhijie Liu, Ying Zhang

**Affiliations:** Department of Rheumatology and Immunology, Clinical Laboratory and Urology, Medical Research Institute, State Key Laboratory of Virology and Biosafety, Frontier Science Center for Immunology and Metabolism, Zhongnan Hospital of Wuhan University, Wuhan University, Wuhan 430071, China; iHuman Institute, ShanghaiTech University, Shanghai, China; RNA Institute, Hubei Key Laboratory of Cell Homeostasis, TaiKang Centre for Life and Medical Sciences, TaiKang Medical School, Wuhan University, Wuhan, 430071,China

## Abstract

The CRISPR–Cas9 system locates genomic targets through gRNA pairing and recognition a protospacer-adjacent motif (PAM). While PAM specificity is typically sequence-determined, we previously found that DNA topology can regulate PAM specificity, relaxing PAMs of various Cas9 effectors and allowing near-PAMless cleavage activity of a type II-C *Alicyclobacillus tengchongensis* Cas9 (AtCas9). However, the structural mechanism underlying this regulation remains unknown. Here, we report cryo-EM structures of AtCas9 bound to B-form DNA or a 340-bp underwound minicircle DNA bearing WT or mutant PAMs. Despite differences in PAM sequences, all three underwound complexes adopt an almost identical architecture that is distinct from the B-form DNA-bound state. On B-form DNA, AtCas9 recognizes the PAM via base-specific hydrogen bonds and steric exclusion, conferring preference for N_4_CNNN and N_4_RNNA (R = A/G). In contrast, underwound minicircle DNA widens the PAM major groove, reduces steric clashes, and enables a compensatory mode involving sequence-independent backbone contacts, explaining near-PAMless cleavage under underwound conformation. These findings uncover a topology-dependent mechanism of PAM recognition and establish a cryo-EM platform using underwound minicircle DNA for structural studies under native-like topological states.

## Introduction

DNA topology refers to the three-dimensional configuration of intertwined DNA duplexes. Owing to the extreme length of genomic DNA and its confinement within compact nuclear spaces, the physical state of DNA, including overwinding (*i*.*e*.,positive supercoiling) and underwinding (*i*.*e*.,negative supercoiling), profoundly influences nearly all nucleic acid processes^1-3^. Despite its significance, how DNA topology regulates protein-DNA interactions remains poorly understood, with no structural studies elucidating the underlying mechanism.

The CRISPR/Cas9 system is a powerful genome-editing platform, in which target recognition relies on both guide RNA (gRNA) complementarity and the presence of a short protospacer adjacent motif (PAM) on the target sequence^4-7^. PAM recognition, mediated by the PAM-interacting (PI) domain of Cas9, acts as a molecular “key” that grants access to the DNA target. While variations in PAM binding models explain diversity in sequence requirements^5,8,9^, structural and mechanistic models have largely been derived from DNA in a single topological state, linear B-form DNA. Beyond sequence constraints, DNA topology, such as underwinding, can modulate Cas9 activity by promoting local DNA unwinding and facilitating R-loop formation^10-12^. However, how DNA topology directly shapes PAM recognition, and whether the PI domain employs a distinct recognition mechanism under topological stress, remains unknown.

Our previous work identified a type II-C thermophilic *Alicyclobacillus tengchongensis* Cas9 (AtCas9), whose PAM specificity is strongly modulated by DNA topology^13^. When DNA adopts an underwound conformation, AtCas9 displays near-PAMless cleavage activity in *E*.*coli*^13^, a property not explained by canonical models of PAM sensing. In addition to its unusual PAM behavior, AtCas9 exhibits optimal nuclease activity at 55 °C and retains robust editing activity in mammalian cells^13,14^, highlighting its potential as a versatile genome editing tool. The unique near-PAMless cleavage activity of AtCas9 makes it an ideal model for studying the impact of DNA topology on CRISPR-based systems. To uncover the molecular basis of this phenomenon, we determined cryogenic electron microscopy (cryo-EM) structures of AtCas9 bound to distinct DNA topoisomers: linear B-form and underwound minicircle DNA. In linear DNA, a loop from the PI domain docks into the major groove of the PAM region, engaging in a base-specific hydrogen bonding and steric exclusion to enforce sequence specificity. Under underwound conditions, the widened major groove relaxes these steric constraints present in B-form DNA, allowing a compensatory recognition mode in which AtCas9 utilizes sequence-independent backbone contacts. These contacts made accessible by DNA topology, enhance protein-DNA interactions and stabilize PAM engagement. These findings uncover a previously unrecognized, DNA topology-regulated mechanism of PAM recognition, demonstrating that DNA conformation, not just sequence, can govern Cas9 specificity. This expands the mechanistic framework of CRISPR-Cas9 targeting and suggests that genome topology is an active regulatory layer influencing editing outcomes.

## Results

### Structures of dAtCas9/sgRNA in complex with B-form DNA bearing WT PAM

*Alicyclobacillus tengchongensis* (AtCas9) belongs to the type II-C Cas9 family and naturally recognizes a permissive PAM sequence of N_4_CNNN or N_4_RNNA (R = A/G) (**Fig. 1a** and **Extended Data Fig. 1a**), when DNA is in canonical B-form^13^. In contrast, underwound DNA topology permits PAMless binding and cleavage by AtCas9 both *in vitro* (**Fig. 1a** and **Extended Data Fig. 1a**) and in *E. coli*^13^. To investigate how DNA topology regulates PAM recognition, we sought to resolve cryo-EM structures of AtCas9-sgRNA complexes bound to DNA substrates with defined topological states. A catalytically inactive AtCas9 variant (dAtCas9, D8A/H617A/N640A) was used to facilitate complex formation (**Supplementary Fig. 1a-b**). To obtain a stable and homogeneous ribonucleoprotein (RNP) complex for structure analysis, we first optimized the sgRNA. Spacer sequences were screened to minimize spacer-dependent interference with sgRNA scaffold folding, as predicted by secondary structure modeling and validated by electrophoretic mobility shift assays (EMSA). Among these, the 21C spacer showed the most favorable profile, strong binding affinity and minimal interference with scaffold architecture (**Extended Data Fig. 1b-c**). Given that sgRNA scaffold influences the stability of RNP assembly and activity^15^, we further modified the scaffold by truncating the repeat:anti-repeat duplex and/or stem-loop 2, and by re-engineering stem-loop 1 into an extended configuration (**Extended Data Fig. 1d**). *In vitro* cleavage assays revealed that several scaffold variants (v1, v2, v5, v6, v8) retained robust catalytic activity, while shortening the spacer to 20-nt diminished cleavage efficiency (**Extended Data Fig. 1e-f**), consistent with our previous in-cell cleavage assays^13^. Based on these results, we selected the 21C-sgRNA-v5, comprising a 24-nt spacer and a 93-nt optimized scaffold (117-nt total), for cryo-EM analysis.

**Figure 1.**
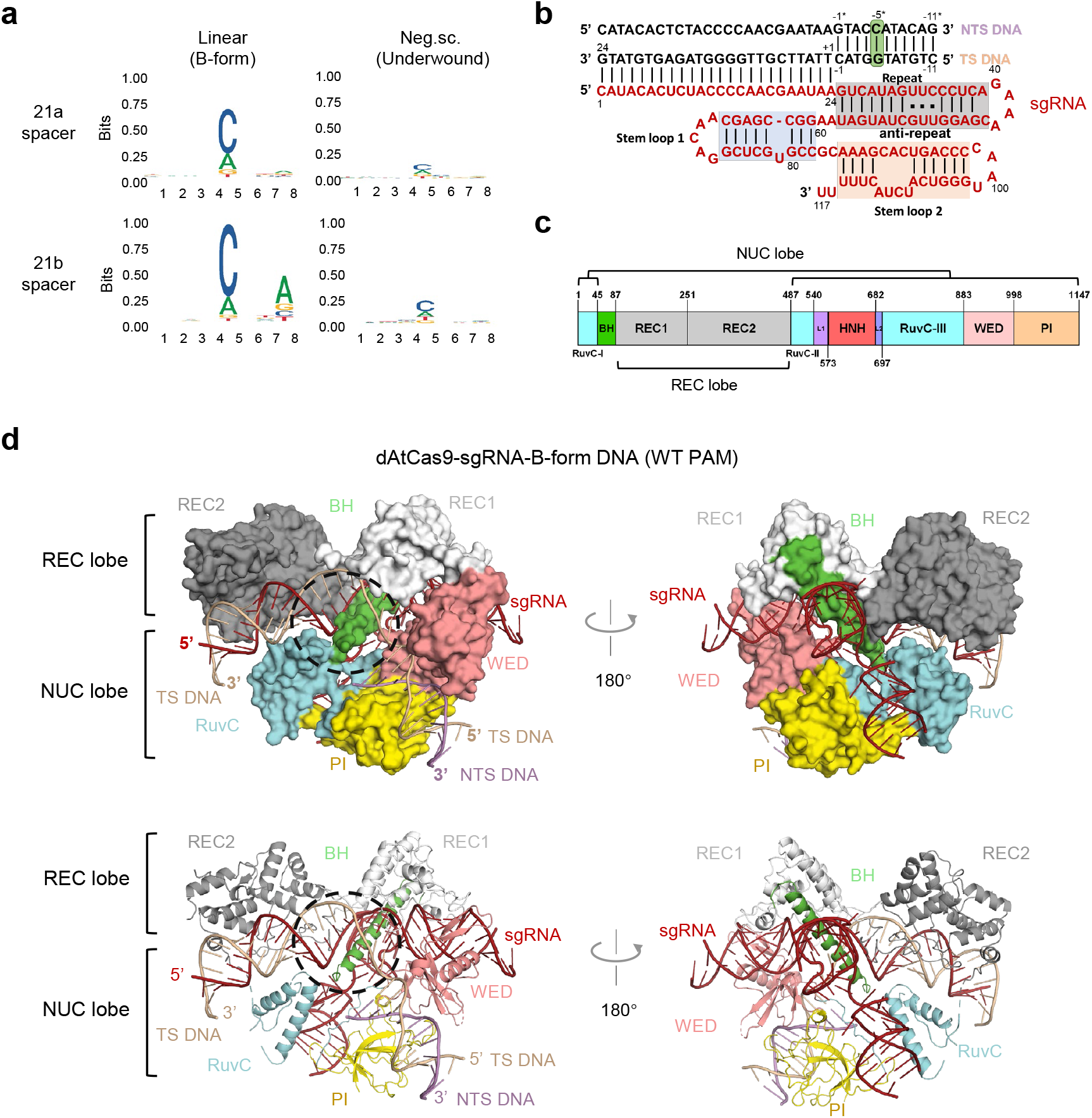
The overall structures of dAtCas9/sgRNA in complex with B-form DNA. **a**, Sequence logos for AtCas9 PAM preference using two spacers 21a and 21b. Neg.sc., negative supercoil. Linear, plasmid DNA were linearized. **b**, Schematic representation of the 35-bp B-form dsDNA bearing WT PAM and 117 nucleotides (nt) sgRNA used for cryo-EM. The fifth cytosine in the WT PAM (5′-N_4_CATA-3′) is highlighted in green. The repeat:anti-repeat region is highlighted in gray. Stem-loops 1 and 2 are highlighted in blue and orange, respectively. **c**, Domain organization of AtCas9. BH, bridge helix; L1, linker 1; L2, linker 2. **d**, Illustration and surface representation of dAtCas9 with sgRNA and a fully complementary linear dsDNA bearing WT PAM, with a 180° rotated view shown on the right. The unresolved HNH domain is indicated by a black dashed circle. Individual AtCas9 domains are colored according to the scheme in (c).

We solved the cryo-EM structure of full-length dAtCas9 (residues 1-1147) in complex with a 117-nt engineered sgRNA and 35-bp linear double-strand DNA (B-form) containing a 5’- N_4_CATA-3’ WT PAM, at 3.15 Å resolution (**Fig. 1b-c, Extended Data Fig. 2a-e** and **Supplementary Table 1**). AtCas9 adopts the canonical bilobed configuration observed in other Cas9 orthologs, comprising a recognition (REC) lobe and a nuclease (NUC) lobe, which are connected by an arginine-rich bridge helix (BH) (**Fig. 1c-d**). The REC1 and REC2 domains are smaller than those of type II-A enzymes, consistent with type II-C orthologs (**Extended Data Fig. 3a-b**), and the NUC lobe contains the RuvC, WED, HNH and PAM-interacting domains. The HNH endonuclease domain, responsible for cleaving the target DNA strand, and its flanking linkers (residues 540–697) are not resolved, consistent with the highly dynamic HNH conformations reported for CjCas9, CdCas9 and GeoCas9^16-18^ (**Fig. 1d** and **Extended Data Fig. 3a**). The target strand (TS) forms a 23-nt heteroduplex with the complementary guide RNA, accompanied by an 11-bp PAM-containing duplex segment that is kinked by ∼71° relative to the R-loop (**Extended Data Fig. 3c**), a geometry consistent with full R-loop formation observed in other Cas9 orthologs^19,20^. The unwound NTS DNA strand is not resolved in the current reconstruction.

The sgRNA, comprising the repeat:anti-repeat duplex followed by two stem-loop structures, is positioned in the positively charged cleft formed by the REC and NUC lobes. The solvent-exposed loop regions (G40-C44, A70-G73, A99-U101 and A111-U117) lack sufficient stability and therefore do not yield interpretable density. AtCas9 makes extensive hydrogen-bond contacts with the sgRNA scaffold (**Extended Data Fig. 4**). The repeat:anti-repeat duplex (G25-U58) adopts an A-form-like conformation. Of its 14-bp paired region, the proximal 12 bp are specifically engaged by the BH, REC1, and WED domains. Consistent with this structural observation, truncating the duplex to a 14-bp complementary region was sufficient to retain full cleavage activity (**Extended Data Fig. 1d-e**). Stem-loop 1 (G61-C82) is recognized by the BH and REC lobes, whereas the stem-loop 2 (A85-U117) primarily interacts with the NUC lobe. The engineered U79 bulge, introduced based on sequence alignment with Nme2Cas9 to enhance activity (V5 variant in **Extended Data Fig.1d-e**), forms several contacts with the BH (Arg76) and REC1 (Gln249 and Arg250), suggesting that increased non-covalent interactions between the stem loop and Cas9 may underlie the observed activity improvement. Together, these features illustrate that AtCas9 relies on a compact yet highly cooperative and conservative set of sgRNA-protein interactions to ensure stable complex formation and efficient cleavage.

### Structures of dAtCas9/sgRNA in complex with underwound minicircle DNA bearing WT PAM or bearing MUT PAMs

Next, we sought to determine the structure of the dAtCas9-sgRNA complex bound to underwound DNA bearing WT or MUT PAMs. Because AtCas9 strongly prefers a cytosine at PAM position 5 and largely rejects thymine at this position (**Extended Data Fig. 1a**), we selected a WT PAM (N_4_CATA) and two T-substituted mutant PAMs (N_4_TATA and N_4_TTGA) for structural analysis. To generate topologically constrained DNA substrates, we adapted the Ligase-Assisted Minicircle Accumulation (LAMA) method^21^ and introduced ethidium bromide (EB) to enable production of underwound topoisomers. In our approach, two PCR-amplified 340-bp fragments were denatured and annealed to form nicked circular DNA, followed by repeated denature-annealing cycles in the presence of thermostable Taq DNA ligase (**Fig. 2a**). During ligation, EB intercalation between base pairs induced DNA unwinding, thereby trapping the desired underwound conformations. Incomplete hybrids and linear fragments re-enter the cycle, leading to the progressive enrichment of covalently closed circular DNA (**Fig. 2a**). By adjusting EB concentration, we systematically tuned the superhelical density (σ) of the 340-bp minicircle from 0 to -0.13 (**Extended Data Fig. 5a**). EMSA confirmed that increasing levels of DNA underwinding enhanced AtCas9 affinity for DNA substrates bearing the mutant PAM (**Extended Data Fig. 5b**). To independently validate the topological state of EB-generated underwound minicircles, we treated the 340-bp minicircles with topoisomerase 1 (TOP1) or gyrase, which respectively relax or introduce negative supercoils in plasmid DNA (**Supplementary Fig. 2a-b**). Although enzymatic conversion was only partial in minicircles, the resulting shifts in topoisomer distributions confirmed the expected changes in σ (**Supplementary Fig. 2c-d**). Using TOP1-relaxed minicircles in AtCas9 cleavage assays, we observed reduced cleavage activity relative to the untreated 12EB substrate (**Extended Data Fig. 5c** and **Supplementary Fig. 3a**), mirroring the effects of EB-induced topology changes and independently verifying the topological status of the EB-generated underwound minicircles. Given that plasmid DNA isolated from *E*.*coli* typically exhibits a superhelical density of -0.06 to -0.07, we selected 12 μg/mL EB to generate a distribution of three topoisomers (σ= -0.06 to -0.13), with the predominant species at -0.09. This condition was used to assemble the dAtCas9-sgRNA-underwound DNA ternary complex for structural analysis.

**Figure 2.**
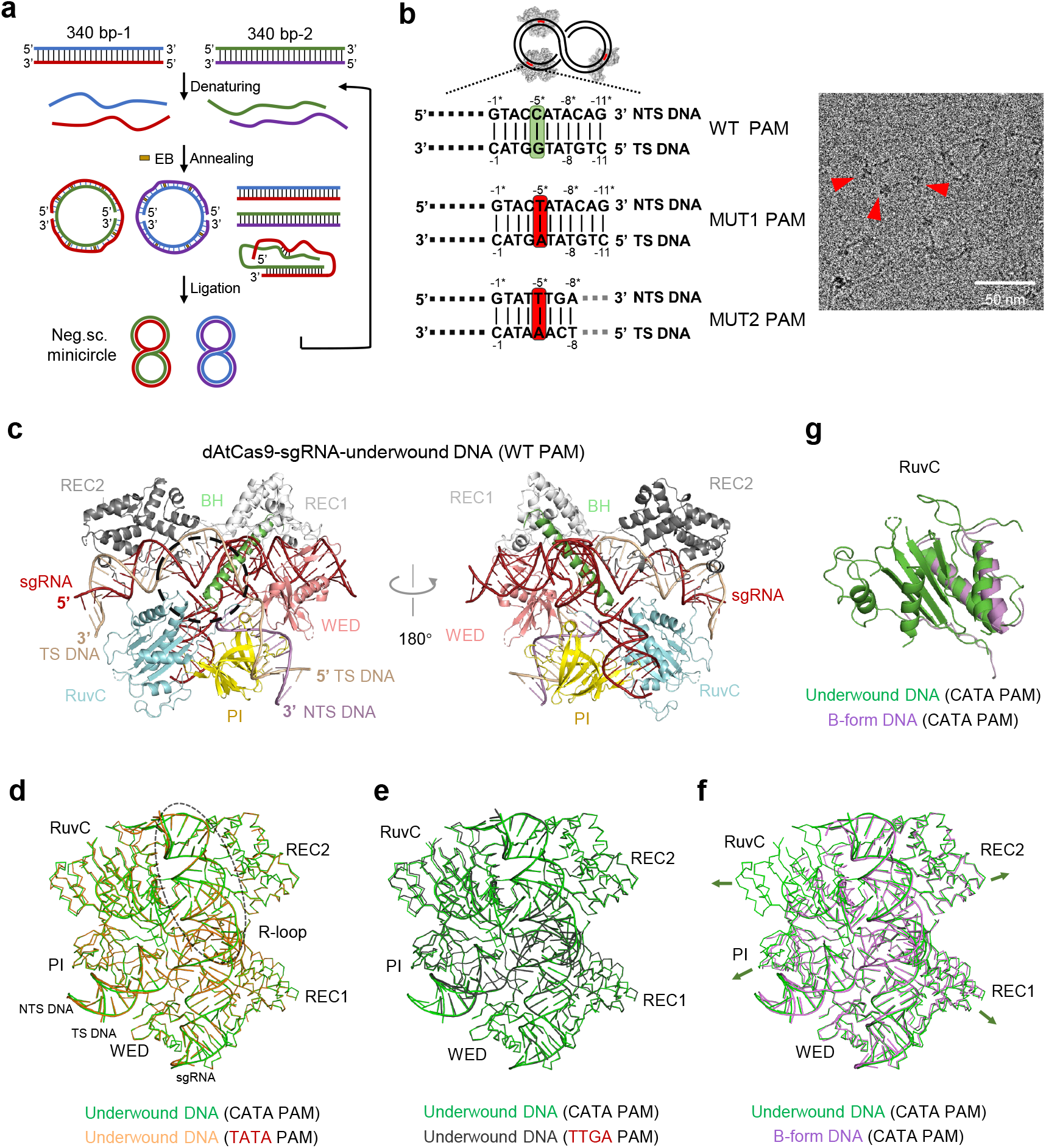
The overall structures of dAtCas9/sgRNA in complex with underwound minicircle DNA bearing WT PAM or MUT PAMs. **a**, Schematic of underwound DNA minicircle generation from linear dsDNA. Two 340-bp linear DNA fragments are denatured and annealed to form nicked circular DNA, partially hybridized linear DNA, or fully hybridized linear duplexes. In the presence of Taq DNA ligase and incorporating the DNA intercalator, ethidium bromide (EB), only nicked circular DNA is ligated to form underwound closed circles, while linear DNA is denatured and recycled in the next cycle. **b**, (Left) Schematic of the dAtCas9-RNP complex bound to three target sites on underwound minicircle DNA, along with the PAM sequence of the target DNA. The fifth cytosine in the WT PAM (5′-N_4_CATA-3′) is highlighted in green, the corresponding thymine in mutant PAM 1 (5′-N_4_TATA-3′) and mutant PAM 2 (5′-N_4_TTGA-3′) is highlighted in red. (Right) Representative raw cryo-EM micrographs of the dAtCas9-sgRNA-underwound minicircle DNA complex with three binding sites. RNP particles bound to minicircle DNA are indicated by red arrows. **c**, Cartoon representation of the dAtCas9-sgRNA-underwound DNA ternary complex bearing the WT PAM. The disordered HNH domain is outlined by a black dashed circle. **d-e**, Structural comparison of AtCas9 bound to underwound DNA containing WT PAM(CATA), MUT1 PAM(TATA), or MUT2 PAM (TTGA). A fully formed R-loop is indicated by black dashed lines. **f**, Structural comparison of AtCas9 bound to B-form DNA and underwound DNA containing the WT PAM. **g**, RuvC domain comparison between the B-form DNA complex and the underwound DNA complex, both containing the WT PAM. NTS, non-target strand; TS, target strand; PAM, protospacer adjacent motif; sgRNA, single guide RNA.

Initial cryo-EM analysis of the dAtCas9-sgRNA complex bound to a 340-bp underwound minicircle containing a single target site revealed severe preferred particle orientation (**Extended Data Fig. 5d**), likely due to the planar distribution of the large circular DNA on the ice layer. This limited angular sampling and hindered high-resolution reconstruction. To resolve this, we redesigned the minicircle to carry three identical target sites, spaced to avoid integer multiples of the DNA helical repeat. This configuration increased the diversity of AtCas9 orientations relative to the DNA ring, thereby improving the angular distribution (**Fig. 2b**). Using this strategy, we collected 5,487 untilted and 2,378 30°-tilted micrographs of dAtCas9/sgRNA bound to underwound minicircle DNA bearing WT PAM (**Fig. 2c** and **Extended Data Fig. 5e-h**). For the MUT1 and MUT2 PAM substrates, we acquired 6,718/2,090 and 7,028/1,449 untilted/tilted micrographs, respectively (**Extended Data Fig. 6**). This enabled high-resolution reconstructions of the active-state dAtCas9-sgRNA-underwound DNA ternary complex at 2.41 Å (WT), 2.65 Å (MUT1) and 2.89 Å (MUT2) (**Fig. 2c-e** and **Supplementary Table 1**). Structurally, the three underwound DNA complexes bearing different PAM sequences (CATA, TATA and TTGA) are highly similar to one another and their protein domain organization is essentially identical (**Fig. 2d-e**). Superposition of the underwound DNA bound complexes shows excellent fit regardless of PAM sequence, indicating a conserved conformation. By contrast, comparison of the WT underwound complex with the B-form DNA-bound complex, which share identical DNA sequences and differ only in topology, reveals a clear difference that the entire protein-DNA assembly in the underwound complex is displaced outward relative to the B-form complex (**Fig. 2f**). This global outward shift appears to be a concerted conformational adjustment that accommodates the altered helical geometry of the underwound DNA. The underwound minicircle complex shows well-resolved density for the RuvC domain (residues 2-22, 28-45, 487-537, 698-737, 752-767, 834-883), whereas only residues 490-498, 503-528, 699-717 and 870-883 are visible in the B-form DNA-bound complex (**Fig. 2g**), indicating that DNA underwinding stabilizes and orders this region. Similar with the dAtCas9-sgRNA-B form DNA complex, the HNH domain remained unresolved, likely reflecting its intrinsic conformational flexibility.

All three underwound DNA structures consistently exhibit a 24-bp RNA-DNA hybrid, compared to 23-bp hybrid observed in the B-form DNA ternary complex (**Extended Data Fig. 7a-c**), indicating that DNA underwinding facilitates R-loop propagation and enable AtCas9 to stabilize a more extended hybrid. Consistently, truncating R-loop to 20 nt reduced cleavage activity for linear and underwound DNA, whereas no discernable differences were observed among hybrids of 21-24 nt (**Extended Data Fig. 7d** and **Supplementary Fig. 3b**), further supporting full R-loop formation in the MUT PAM under underwound conditions. The phosphate-lock loop (Glu886-Thr887), which stabilizes the sharp kink in the target strand upstream of the PAM, is properly formed in the underwound structures (**Extended Data Fig. 7e**). The correct positioning of this loop further indicates that the canonical early steps of R-loop establishment are preserved.

At the level of sgRNA recognition, AtCas9 engages nearly identical numbers and types of non-covalent interactions with the sgRNA scaffold between underwound complexes of WT or MUT PAM, consistent with their overall structural similarity (**Extended Data Fig. 8-9**). While most sgRNA-protein contacts are shared between the underwound and B-form structures, several interaction differences are evident, showing five B-form unique contacts and eight underwound unique contacts. Notably, seven of the eight underwound-specific contacts originate from domains that undergo topology-induced movements, REC1, WED and RuvC, with one additional residue from the BH domain. For instance, in the underwound structure, the +6 position of the target strand is displaced outward, prompting an adjacent REC1 residue, Ser137, to shift slightly and reduce its distance to 6C(OP1) from 4.4 Å to 3.3 Å, forming an underwound-specific hydrogen bond (**Extended Fig. 10a**). A similar effect occurs in the WED domain, where Arg973 moves closer to sgRNA G51 to establish a topology-specific contact (**Extended Fig. 10b**). Conversely, conformationally adaptive shifts can increase local distances and abolish hydrogen bonding, as exemplified by the loss of the B-form contact involving Leu55 in BH domain (**Extended Fig. 10b**).

### Structural divergence of the PAM duplex

Next, we compared the sgRNA: DNA heteroduplex between the B-form and underwound complexes, which were nearly identical, showing high structural concordance in the phosphate backbone, deoxyribose positioning, and base stacking (**Fig. 3a-b**). In contrast, marked differences were observed in the PAM duplex region, where both the phosphate backbone and base pairs exhibited pronounced deviations between the B-form and underwound conformations (**Fig. 3b**). In the underwound DNA complex, the major groove was notably widened, with a width of 21.6 Å or 21.2 Å for two different PAMs respectively, compared to 19.3 Å in the B-form complex (**Fig. 3c**). Helical parameters analysis using 3DNA further supported this difference: the underwound DNA exhibited a reduced base-step twist and a reduced rise per base pair **(Extended Data Fig. 10c**).

**Figure 3.**
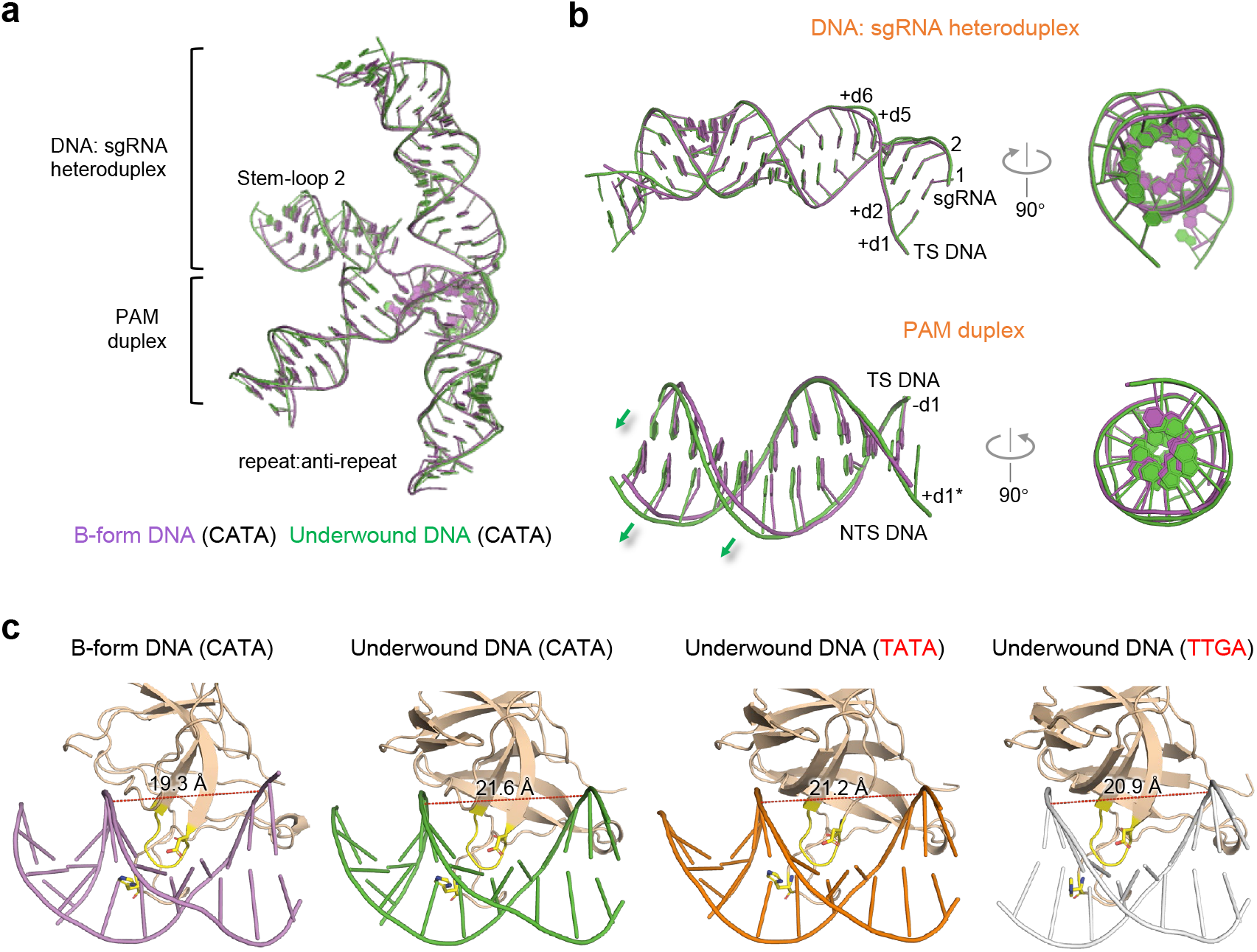
Structural divergence of the PAM duplex **a**, Structural comparison of sgRNA-target DNA complexes of B-form DNA and underwound DNA. **b**, Enlarged side views of the superimposed DNA:sgRNA heteroduplex and PAM duplex, with a 90° rotated view shown on the right. The expansion direction of the underwound DNA relative to B-form DNA is indicated by green arrows. **c**, Comparison of DNA phosphate backbone structures. The geometry of major groove is measured and highlighted as red dashed lines. In the PI domain, loops inserted into the major groove are shown in yellow, with the key base-specific interacting residue (D1089 and H1034) displayed in stick representation.

To quantitatively assess these topological effects, we modeled a 340-bp minicircle DNA with a superhelical density of σ = –0.09 or –0.06, assuming that torsional stress is fully absorbed through underwinding without writhe formation. In the absence of Cas9, the minicircle adopted a helical pitch of ∼11.5 bp/turn or 11.2 bp/turn and a base-step twist of 31.3°or 32.1°, respectively (**Extended Data Fig. 10c**). For comparison, canonical B-form DNA has a helical pitch of ∼10.5 bp/turn and a base-step twist of 34.3°. In our minicircle DNA system, torsional stress is constrained and the single minicircle contains three identical Cas9-binding sites. Thus, binding of Cas9 at one site would be expected to influence the topology of the remaining two sites. We reason that this redistribution of stress partially influences the PAM duplex region, as the base-step twist observed in the cryo-EM structure was higher than the calculated value, indicating a compensatory increase in twist to offset stress from R-loop formation. Consequently, the PAM duplex retains its underwound conformation, albeit partially relaxed, and remains clearly distinct from the B-form DNA complexes of other Cas9 orthologs (**Extended Data Fig. 10c**), further validating the physiological relevance of the cryo-EM structure.

### AtCas9 PAM recognition is intrinsically sensitive to the steric environment at PAM position 5

Structural analysis revealed that the PAM region is stabilized by a loop from the PI domain of AtCas9 docking into its major groove (**Fig. 3c**). Our previous work defined the preferred PAM for AtCas9 as N_4_CNNN or N_4_RNNA^13^. Consistent with this, the carboxyl group of Asp1089 forms a hydrogen bond (3.3 Å) with the 4-amino group of -dC5* on the NTS, while the epsilon-amino group of Lys1107 forms a hydrogen bond (3.4 Å) with the O6 atom of -dG5 on the TS, jointly specifying PAM recognition (**Fig. 4a**). In addition, His1034 from AtCas9 engages position 8 on the TS via hydrogen bonding to -dT8, corresponding to an A on the NTS (**Fig. 4a**). When a substitution at position 5 from A to T was modeled using PyMOL, the resulting structure revealed a steric clash between the 5-methyl group of thymine and the side chain of Asp1089 (**Fig. 4a**), providing a structural rationale for the exclusion of thymine at this position. Supporting this model, replacing T with uracil, which lacks the methyl group, markedly enhances DNA cleavage, whereas introducing a C5-methyl group strongly inhibits AtCas9 activity (**Fig. 4b** and **Supplementary Fig. 3c**). These strongly suggest that in B-form topology, alteration of the methyl group at PAM position 5 can substantially influence AtCas9 binding and cleavage. Together, these data explain AtCas9’s preference for cytosine at PAM position 5 and adenine at position 8 in B-form DNA.

**Figure 4.**
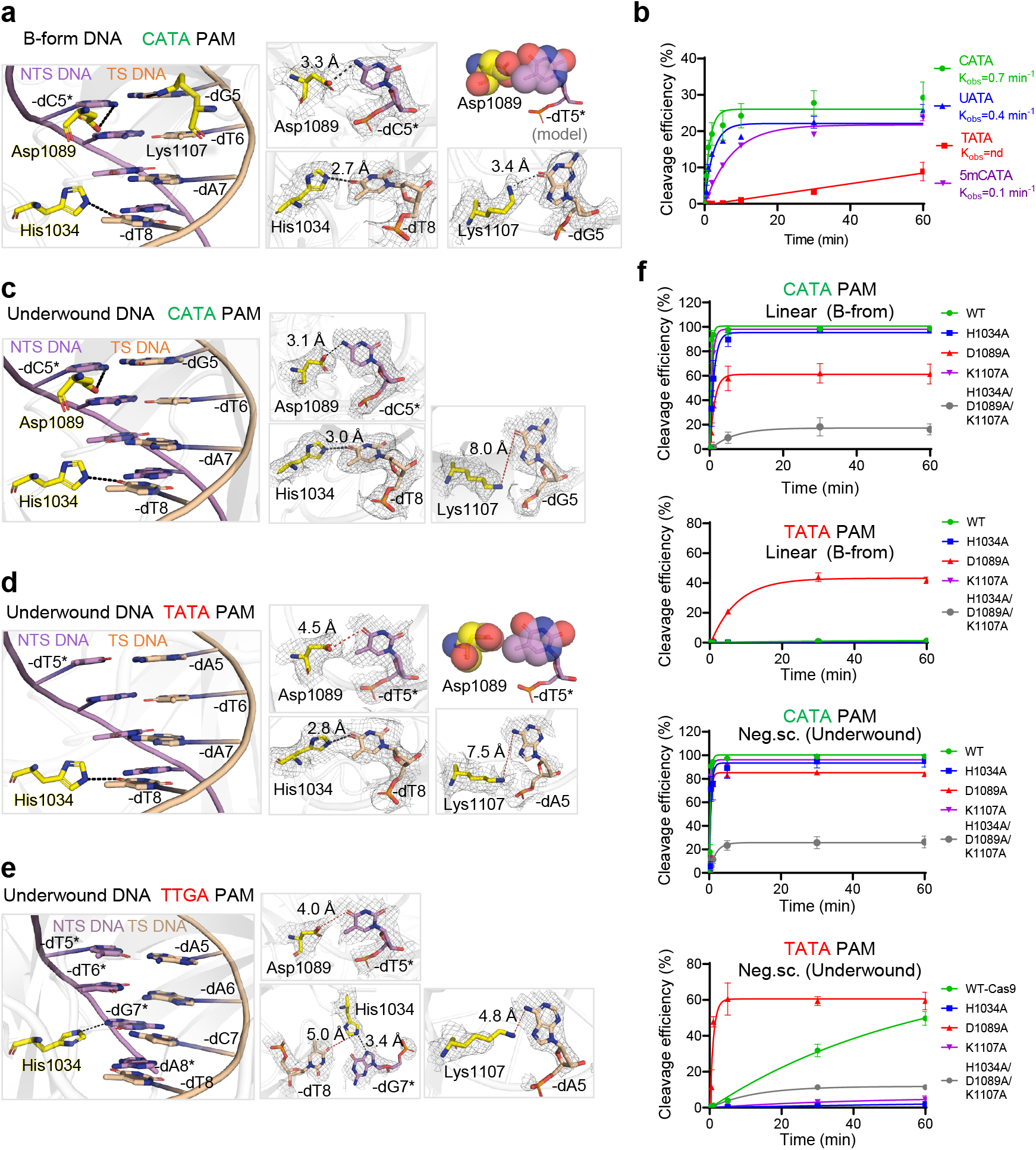
PAM recognition mechanism of B-form DNA and underwound DNA. **a**, Enlarged view of base-specific interactions between the PI domain and the PAM duplex in the B-form DNA ternary complex (Left). Close-up views of the specific hydrogen-bond interactions of Asp1089, His1034 and Lys1107 with the corresponding bases are shown, with omitted EM density for clarity and the residues represented in stick model. Asp1089 is shown in sphere representation together with the modeled mutant –dT5* (generated in PyMOL) (Right). Hydrogen bonds are indicated by gray dashed lines. **b**, *In vitro* cleavage kinetics of AtCas9 on B-form substrates carrying PAM variants differing at position 5 (C, T, U or 5mC). 5mC denotes a methylated cytosine (also see Supplementary Fig. 3c). **c-e**, Enlarged view of base-specific interactions between the PI domain and the PAM duplex in underwound DNA containing the WT PAM (**c**), MUT1 PAM (**d**) or MUT2 PAM (**e**). Close-up views of the hydrogen-bond interactions (gray dashed line) and atomic distance (red dashed line) between Asp1089, His1034, Lys1107, and their corresponding bases are shown. **f**, *In vitro* cleavage kinetics of AtCas9 bearing indicated mutations on WT PAM (CATA) and mutant PAM (TATA) under different DNA topologies (also see Supplementary Fig. 3d). Data are represented as mean ± SD and fitted with non-linear regression.

### Differential recognition of canonical and mutant PAMs in underwound DNA topology

In the underwound WT PAM complex, the PAM-proximal geometry around position 5 resembles that of the B-form DNA complex: Asp1089 maintains a hydrogen bond (3.1 Å) with dC5* on the NTS (**Fig. 4c**). By contrast, in the two mutant PAM complexes (N_4_TATA and N_4_TTGA), underwound DNA allows AtCas9 to tolerate thymine at position 5, a base that is normally disfavored in B-form DNA. Although no base-specific contacts are observed between the PI domain and the -dT5*, structural comparison revealed that the distance between Asp1089 and -dT5* increases to 4.5 Å and ∼4.0 Å in the two underwound DNA structures (**Fig. 4d-e**). This expanded distance eliminates the steric clash between the thymine 5-methyl group and Asp1089, a clash that underlies thymine exclusion in the B-form context. These changes are consistent with the altered local geometry and widened major groove characteristic of underwound DNA (**Fig. 3c**), which collectively shift the PAM-adjacent duplex outward. This topologically induced relaxation reduces the steric penalty at position 5, improves PAM exposure, and makes the site more permissive for Cas9 loading. Together, these features provide a structural explanation for the markedly relaxed specificity toward N_4_TNNN PAMs when DNA adopts an underwound topology.

In all three underwound complexes, the side chain of Lys1107 adopts a swung-out conformation that increases its distance from the –d5 position, thereby abolishing its interaction with PAM duplex and exhibiting a conformation-dependent mode of recognition (**Fig. 4c-e**). Furthermore, His1034 consistently engages the O4 atom of –dT8 on the TS, exhibiting only a subtle positional displacement between the B-form and underwound states (CATA and TATA PAMs) (**Fig. 4a** and **Fig. 4c-d**). In the TTGA PAM context, His1034 no longer contacts –dT8, but instead repositions to interact with -dG7* on the NTS via a new hydrogen bond (**Fig. 4e**). Thus, while His1034 preferentially interacts with position 8 on the target strand, its contacts remain sequence-dependent.

In the B-form complex, PAM specificity is established by three base-specific contacts together with steric exclusion between Asp1089 and thymine at PAM position 5. Alanine substitution of any of these residues reduced cleavage activity to varying degrees (**Fig. 4f** and **Supplementary Fig. 3d**), with D1089A causing the most pronounced defect, consistent with its dominant contribution to PAM recognition. The triple mutant fully abolished activity. When encountering mutant PAM carrying thymine at position 5, shortening the Asp1089 side chain, as in D1089A, partially relieves this steric constraint and improves tolerance for thymine. Accordingly, D1089A increased cleavage activity of MUT PAM (N_4_TNNA) substrates, with a greater effect on linear DNA than underwound DNA (**Fig. 4f**), likely because underwinding already mitigates this steric barrier and thus partially reduces the benefit of the mutation. In contrast, mutation of His1034 or Lys1107 only mildly affected cleavage of N_4_CNNA substrates but substantially impaired activity on N_4_TNNA MUT PAMs (**Fig. 4f**). These results indicate that although DNA topology can partially compensate for weakened PAM recognition, the resulting affinity remains intrinsically limited and can be compromised when additional stabilizing interactions are lost, thereby diminishing the topology-enabled cleavage activity.

### Sequence-independent contacts contribute to PAM recognition in underwound DNA

In resolved Cas9-DNA structures, sequence-specific PAM recognition provides an anchor point that allows Cas9 to bend the DNA into a kinked conformation, thereby facilitating RNA-DNA base-pairing to initiate DNA unwinding^19^. Although underwound DNA introduces negative torsional stress that energetically favors DNA unwinding, we hypothesize that non-specific interactions may also contribute to AtCas9’s ability to initiate DNA bending. In support, the PI domain makes extensive contacts with the phosphate backbone of DNA, particularly within the PAM region (**Fig. 5a**). Due to the distinct DNA topology, the underwound complexes exhibit a set of PAM-proximal, non-sequence-specific backbone interactions that are identical across all PAM variants and arise solely from the underwound conformation. For example, Lys980 and Lys917 positioned on the periphery of the DNA duplex, form hydrogen bonds with the DNA backbone in the underwound structures (**Fig. 5a-c**), but are absent in the B-form complex (**Fig. 5c**). Consistent with this, alanine substitution of these residues selectively reduced cleavage activity on underwound substrates while leaving B-form DNA cleavage unaffected (**Fig. 5e** and **Supplementary Fig. 3e**), indicating that these contacts stabilize the underwound PAM geometry.

**Figure 5.**
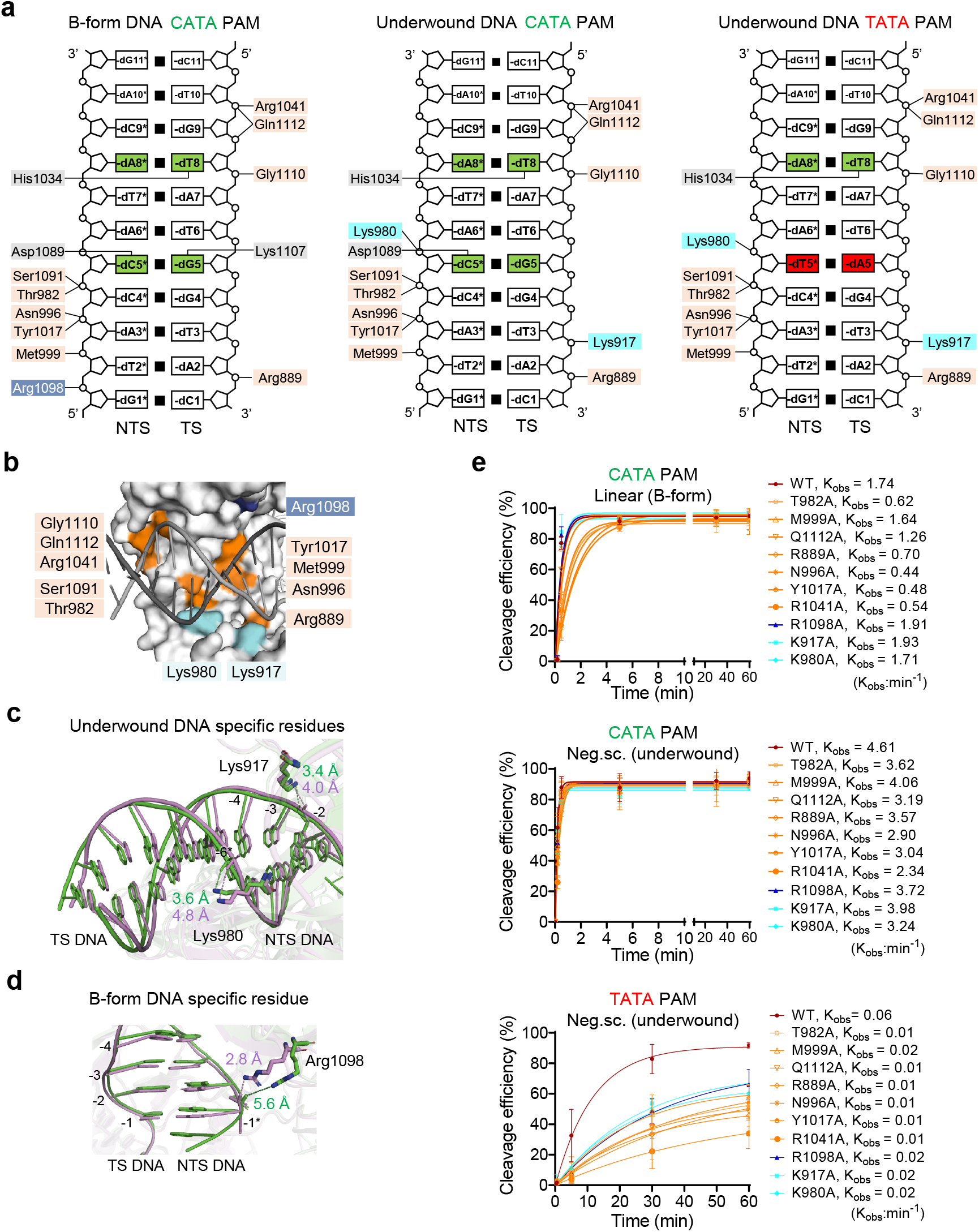
Sequence-independent contacts between the PI domain and PAM duplex. **a**, Schematics of the PAM duplex contacts by AtCas9 in B-form DNA and underwound DNA containing either the WT PAM or MUT1 PAM. Hydrogen bonds are depicted by black lines. Cyan highlights underwound-specific contacts; Blue, B-form-specific contacts; Brown, shared contacts between the two topologies. **b**, Close-up view of PI domain and its backbone interactions with the PAM duplex. **c**, Superposition of the underwound-specific interacting residues in B-form (violet) and underwound DNA (green) complexes containing the WT PAM. Atomic distances are indicated by dashed lines, respectively. **d**, Superposition of the B-form-specific interacting residues in B-form (violet) and underwound DNA (green) complexes containing the WT PAM. Atomic distances are indicated by gray and green dashed lines, respectively. **e**, Kinetic analysis of cleavage efficiency of AtCas9 bearing the indicated mutations on WT or mutant PAM substrates under different DNA topologies. WT, wild type. Cyan, underwound-specific contacts; Blue, B-form-specific contacts; Brown, shared contacts between the two topologies. Data are represented as mean ± SD and fitted with non-linear regression.

In addition to these topology-specific interactions, we identified nine non-base-specific hydrogen bonds that are present across all topological states (**Fig. 5b**). Disrupting these shared interactions markedly decreased activity on MUT-PAM substrates but caused only minor reductions on WT PAM, suggesting that these non-specific contacts become particularly important when base-specific recognition is weakened. For WT PAM, strong base-specific interactions appear sufficient to compensate for the loss of these backbone contacts, consistent with the modest impact on cleavage (**Fig. 5e** and **Supplementary Fig. 3e**). Finally, Arg1098 forms a DNA-backbone interaction exclusively in the B-form complex (**Fig. 5d**); however, the R1098A mutation produced minimal effect on B-form cleavage, indicating that this interaction is not functionally essential (**Fig. 5e** and **Supplementary Fig. 3e**). Together, these findings show that while base-specific interactions dominate recognition in the presence of a cognate PAM, backbone-mediated contacts become critical under conditions of weakened PAM specificity. The additional topology-specific interactions observed in the underwound complexes further illustrate how DNA topology can buffer reduced PAM recognition and facilitate productive Cas9 loading.

### Comparison of PAM recognition across Cas orthologues

Loop-mediated docking mechanism is conserved across multiple type II-C Cas9 orthologs, including Nme2Cas9^22^, GeoCas9^18^, CjCas9^16^, and ThermoCas9^23^, all of which exhibit major groove docking via loops containing base-specific contacts (**Extended Data Fig. 11a-b**). In case of CjCas9, which recognizes an N_3_VRYM PAM, the preference at the fourth position is likewise governed by steric exclusion. Specifically, the methyl group of dT4* would clash with the side chain of Thr913 in the docking loop, thereby favoring V (A/G/C) at this position (**Extended Data Fig. 11a**).

AtCas9 shares high sequence similarity to its homology Nme2Cas9, the latter of which recognizes an N_4_CC PAM (**Extended Data Fig. 11b**). In both nucleases, a conserved aspartate within the docking loop inserts into the major groove to recognize the cytosine at PAM position 5 on the NTS. In Nme2Cas9, an adjacent arginine additionally engages PAM position 6 on the TS together with Asp1028, conferring CC-specific recognition (**Extended Data Fig. 11b**). In AtCas9, this position-6 recognition is absent, contributing to its permissive PAM signature.

In comparison, the type II-A orthologs SpCas9 employs a mechanistically distinct strategy. Its cognate loop does not insert into the major groove; instead, PAM recognition is achieved through the two arginine side chains that contact the GG dinucleotide^5^ (**Extended Data Fig. 11c**). In the PAM-relaxed variant SpRY, these base-specific interactions are removed and replaced with clusters of substitutions that strengthen electrostatic interactions with the PAM-proximal backbone (**Extended Data Fig. 11d**), thereby stabilizing the R-loop independently of PAM identity^24,25^. Despite their different molecular strategies, AtCas9 (under underwound DNA) and SpRY converge on a shared functional principle: reduced dependence on canonical base-specific PAM recognition and increased reliance on non-base-specific interactions. Underwound DNA remodels the AtCas9 interface to lessen PAM-specific constraints while promoting topology-induced interactions, whereas SpRY achieves PAM relaxation through engineered substitution of PAM-interacting residues to reinforce non-specific contacts with the DNA backbone.

## Discussion

In this study, we report cryo-EM structures of AtCas9 bound to B-form and underwound minicircle DNA bearing WT or MUT PAMs, revealing a previously unrecognized, topology-dependent mode of PAM recognition and a structural mechanism for its near-PAMless activity under underwound topology. A key feature of this mechanism is the loop-docking model of the PI domain with the DNA major groove, which is critical for PAM engagement. In the B-form DNA complex, AtCas9 relies on base-specific hydrogen bonding and steric exclusion to establish a defined PAM preference. In contrast, underwound DNA relaxes these steric constraints and widens the major groove, facilitating PI domain docking. This loop-mediated docking mechanism is conserved among type II-C Cas9 orthologs, and its sensitivity to DNA topology may explain both the relaxed PAM requirements of many type II-C enzymes and the propensity of underwound DNA to enable near-PAMless cleavage. Such relaxed PAM recognition is exemplified by CjCas9, which tolerates NNNVRYM PAMs, and the engineered iGeoCas9, which recognizes N_4_CNNN PAMs^16,18^, highlighting how loop docking can accommodate diverse PAM sequences in type II-C nucleases.

To compensate for limited base-specific interactions, AtCas9 employs extensive sequence-independent contacts, including multiple hydrogen bonds with the phosphate backbones. In addition to these DNA-mediated contacts, the protein forms numerous structure-dependent, non-sequence-specific interactions with the sgRNA scaffold, collectively stabilizing the RNP complex and facilitating target engagement even when PAM recognition is weakened. This highlights a mechanistic principle by which backbone interactions can substitute for canonical base-specific recognition to expand PAM compatibility. A similar strategy underlies the SpRY variant of SpCas9, which achieves near-PAMless activity by strengthening backbone engagement while reducing base-specific PAM contacts^24,25^. Building on this principle, future engineering efforts could further relax the PAM constraints of AtCas9 by strategically diminishing base-specific interactions and reinforcing sequence-independent contacts.

Underwound DNA naturally harbors loosely interwound strands, making it more flexible and easier to bend and unwind. Such topological states profoundly influence the activity of DNA-binding proteins by altering DNA geometry, flexibility, and unwinding kinetics^3^. For example, negative supercoiling destabilizes base stacking, promoting transcription initiation^26^, and enhances DNA bending to facilitate topology-dependent recognition by Factor for Inversion Stimulation (FIS) through adjacent major grooves^27^. Many proteins, such as type I and type II topoisomerases, have evolved to sense and respond to specific topoisomers to maintain topological homeostasis^28^. Similarly, following PAM recognition, Cas9 induces PAM-proximal DNA bending and twisting to facilitate interrogation of the target DNA^19^. We speculate that the intrinsic bendability of underwound DNA lowers the energetic barrier for DNA deformation, potentially accelerating the target search process. Together while underwound DNA reduce the energetic barrier for DNA unwinding and facilitate R-loop propagation, this increased flexibility enhances on-target cleavage efficiency while may simultaneously compromise specificity. Indeed, our previous work and that of others have shown that underwound DNA enhances the off-target activity of Cas9^12,29^.

Despite the importance of DNA topology, structural studies of topology-sensitive proteins have been constrained by technical limitations. Native plasmids are too large for high-resolution cryo-EM, while *in vitro* methods to generate defined topoisomers are inefficient and offer limited control. As a result, most studies rely on linear or partially duplexed DNA, potentially missing key topological insights. To address this, we developed a cryo-EM platform incorporating covalently closed, underwound minicircle DNA with tunable superhelical density and customizable sequence. Using Cas9 as a model, we demonstrate that this system enables structural and biochemical dissection of protein-DNA interactions under native-like topological states. This platform provides a broadly applicable strategy and offers a generalizable strategy for elucidating topology-dependent mechanisms of DNA recognition and regulation.

## Author contributions

M.D. and Y.Z. conceptualized the project and analyzed the data. M.D. purified Cas9, performed in vitro cleavage assays, prepared and collected cryo-EM data with the help of L.Z. and B.M.. B.M. and Z.L. provided critical guidance on cryo-EM experiments. L.W. and B.M. performed the electron microscopy data processing and structure determination. X.T. and D.H. helped with data analysis. H.Y. provided helpful suggestions. M.D. and Y.Z. wrote the manuscript with input from all co-authors.

## Acknowledgments

We thank Dr. Ping Yin (Huazhong Agricultural University) for helpful discussions. This work is kindly supported by Noncommunicable Chronic Diseases-National Science and Technology Major Project (2023ZD0500600), the National Key R&D Program of China (2022YFF1002801 to Y.Z., 2024YFA0917303 to H.Y.,), the National Natural Science Foundation of China (82450105 and 82525033 to Y.Z.; 32501143 to M.D., T2425015 to H.Y.), Ministry of Agriculture and Rural Affairs of China, the Fundamental Research Funds for the Central Universities (2042022dx0003, 2042022kf1190), Wuhan Municipal Key Research and Development Program (2025020602030111), the Postdoctor Project of Hubei Province (2025HBBSHCXB029). We thank the cryoEM Core Facility of Wuhan University and the Bio-Electron Microscopy Facility of ShanghaiTech University for cryo-EM data collection, and the core facility of Medical Research Institute at Wuhan University for their technical supports.

## Materials and Methods

### AtCas9 expression and purification

The plasmid encoding recombinant AtCas9 with a C-terminal 6×His tag was transformed into *E. coli* BL21 (DE3) competent cells. A single colony was inoculated and grown overnight at 37 °C with shaking at 220 rpm. The overnight culture was diluted 1:100 into fresh LB medium and grown at 37 °C until the OD_600_ reached ∼0.6. Protein expression was induced by adding 0.2 mM isopropyl β-D-1-thiogalactopyranoside (IPTG), followed by incubation at 18 °C with shaking at 180 rpm for 18 h. Cells were harvested and resuspended in Buffer A (20 mM Tris-HCl, pH 7.4, 500 mM NaCl) at a ratio of 1:10 (v/v), and then lysed using a high-pressure homogenizer at 4 °C and clarified by centrifugation at 12,000 g for 20 min at 4 °C. The supernatant was filtered and loaded onto a HisTrap HP column (Cytiva) pre-equilibrated with Buffer A. After sample loading, the column was sequentially washed with Buffer A containing 10 mM, 25 mM, and 50 mM imidazole for 5 column volumes (CV) each to remove non-specifically bound proteins. The target protein was eluted with a linear gradient of 50–500 mM imidazole over 10 CV. Eluted fractions were analyzed by SDS-PAGE, and peak fractions were pooled and concentrated to ∼1 mL using a 50 kDa molecular weight cutoff centrifugal concentrator. The protein sample was diluted with low-salt buffer (20 mM Tris-HCl, pH 7.4, 300 mM NaCl) to adjust the salt concentration and loaded onto a HiTrap Heparin HP column (Cytiva) pre-equilibrated with the same buffer. Bound protein was eluted using a linear gradient of 0–100% high-salt buffer (20 mM Tris-HCl, pH 7.4, 1 M NaCl) over 5 CV. Eluted fractions containing the target protein were identified by SDS-PAGE, pooled, and concentrated as described above. Final purification was performed by size-exclusion chromatography using a HiLoad 16/600 Superdex 200 pg column (Cytiva) equilibrated in Buffer B2 (20 mM Tris-HCl, pH 7.4, 300 mM NaCl). Fractions corresponding to the major peak were analyzed by SDS-PAGE, pooled, concentrated to ∼10 mg/mL, aliquoted, flash-frozen in liquid nitrogen, and stored at – 80 °C until further use. The dAtCas9 prokaryotic expression sequence used in this study is listed in Supplementary Dataset 1.

### sgRNA preparation

The sgRNAs were transcribed *in vitro* using T7 RNA polymerase in a 50 μL transcription reaction containing 5 mM DTT, 2 mM each rNTP, 0.1 U pyrophosphatase, 50 U RNase inhibitor, and 500 ng DNA template in transcription buffer (40 mM Tris-HCl, pH 8.0, 25 mM NaCl, 8 mM MgCl_2_, 2 mM spermidine trihydrochloride). Transcription was performed at 37 °C for 2 h, followed by RQ1 DNase (Promega) treatment to remove the DNA template. sgRNAs were purified using the Monarch RNA Cleanup Kit (New England Biolabs). The sgRNA sequences used in this study are listed in Supplementary Dataset 1.

### Electrophoretic mobility shift assay (EMSA)

For the linear dsDNA binding assay, two complementary oligonucleotides (TS and NTS strands) were synthesized (GenScript), with a 5′ FAM label on the TS strand. The oligonucleotides were annealed to generate dsDNA. dAtCas9 and sgRNA were mixed at 1:2 molar ratio in binding buffer containing 10 mM KCl, 20 mM HEPES (pH 7.9), 10 mM MgCl_2_, 0.5 mM DTT, 0.1 mM EDTA. The RNP complex was incubated at room temperature for 10 min, then diluted to the desired concentrations. Subsequently, 5 nM of the dsDNA substrate was added, and the mixture was incubated at 55 °C for 30 minutes. Samples were resolved on 4% native TBE-PAGE. The oligonucleotide sequences used in this study are listed in Supplementary Dataset 1.

For the binding assay with underwound minicircle DNA, circularized DNA was first labeled with Cy5 using the Label IT® Tracker Intracellular Nucleic Acid Localization Kit according to the manufacturer’s protocol (Mirus Bio). Cy5-labeled minicircle DNA (5 nM), containing a mutated PAM sequence (N_4_TTGA), was incubated with the dAtCas9 RNP complex at the indicated concentrations. Binding reactions were resolved on 0.8% agarose gel in sodium boric acid buffer (8.6 mM sodium borate, 45 mM boric acid, pH 8.3) for 3 h at 60 V at 4℃. Gels were imaged by Bio-Rad ChemiDoc MP imager (Bio-Rad).

### *In vitro* cleavage assay

To prepare DNA substrates, oligonucleotides containing the target sequences with various PAMs were synthesized (Sango Biotech) and inserted into the pCE2 plasmid via TA cloning (Vazyme). For linear DNA substrates, supercoiled plasmids were linearized with *NcoI* (New England Biolabs). Purified AtCas9 (100 nM) and sgRNA (200 nM) were first pre-complexed in Buffer 16 containing 10 mM KCl, 20 mM HEPES (pH 7.9), 10 mM MgCl_2_, 0.5 mM DTT, 0.1 mM EDTA at room temperature for 10 min. DNA substrates (negative supercoiled plasmids and linearized plasmids) were then added to a final concentration of 2 nM, and the reactions were performed at either 37 °C or 55 °C for the indicated time. The cleavage reactions were terminated by adding 1 μL of proteinase K (Thermo Fisher Scientific) and incubating at 55 °C for 10 min. Cleavage products were resolved by electrophoresis on a 0.8% TAE agarose gel and visualized by ethidium bromide (EB) staining. For cleavage assays using fluorescently labelled oligonucleotides, AtCas9–sgRNA ribonucleoprotein (RNP) complexes were incubated with DNA substrates at a ratio of 2:1 in buffer 16 at 37 °C for the indicated times. For cleavage assays using 340-bp minicircle DNA, the same RNP:DNA ratio (2:1) was applied, and reactions were performed in buffer 16 at 55 °C for the indicated times. The DNA sequences used for the *in vitro* cleavage assays are listed in Supplementary Dataset 1.

### Determination of superhelical density of 340-bp minicircles

Superhelical density (σ) of 340-bp DNA minicircles was determined by resolving topoisomers on native polyacrylamide gels. Minicircle samples were electrophoresed alongside a relaxed topoisomer control (0EB control), which defines the relaxed (ΔLk = 0) topoisomer. Each band migrating faster than the relaxed species corresponds to an additional unit of negative linking number (ΔLk = –1 per band)^30^. The relaxed linking number (Lk_0_) for a 340-bp minicircle was calculated using 10.5 bp per turn (Lk_0_ ≈ 32–33). Superhelical density was then calculated as:

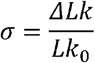

This approach allows assignment of σ values for each topoisomer based on band position relative to the relaxed standard.

### Estimation of structural parameters in topologically constrained DNA

The helical pitch of underwound minicircle DNA was computed based on the relationship between superhelical density (σ), linking number (Lk), and base pair length. For a closed-circular DNA of 340 bp (L = 340) with a defined superhelical density (σ = -0.09), the relaxed linking number (Lk_0_) was estimated using a canonical helical repeat of 10.5 bp/turn:

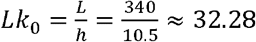

The change in linking number (ΔLk) was calculated from the superhelical density:

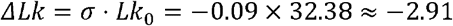

The resulting linking number was:

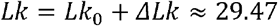

Twist (bp/turn) was computed as:

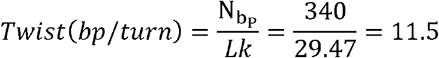

Local base-pair step twist was computed as:

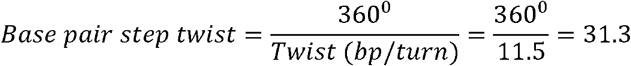

### Substrate B-form DNA preparation

Single-stranded target and non-target DNAs (sequences provided in Supplementary Dataset 1) were synthesized by Sangon Biotech and purified by HPLC. Oligonucleotides were resuspended in nuclease-free water and annealed in the presence of annealing buffer (10 mM HEPES, pH 7.5, 20 mM NaCl, 1 mM EDTA). Target and non-target strands were mixed at a molar ratio of 1:3 in a final volume of 100 μL. Annealing was performed using a thermocycler with the following temperature program: 95 °C for 3 min, followed by gradual cooling at a rate of 0.1 °C/s to 25 °C, and held at 12 °C thereafter.

### Underwound minicircle DNA preparation

Two 340 bp dsDNA fragments (designated 340-A and 340-B; sequences provided in Supplementary Dataset 1) were designed to generate negatively supercoiled circular DNA. Both fragments were PCR-amplified using primers containing 5′ phosphorylation (sequences provided in Supplementary Data set 1) and purified using a PCR purification kit (Magen). For DNA circularization, 400 ng of each fragment was mixed in a 100 μL reaction containing 1 × Taq DNA ligase buffer and the indicated concentrations of EB. The mixture was denatured at 95 °C for 3 min and immediately cooled on ice for 3 min. Taq DNA ligase (80 U, ABclonal) was then added, followed by thermal cycling consisting of 65 °C for 30 min, 95 °C for 20 s, 4 °C for 2 min, and 65 °C for 30 min, repeated for a total of seven cycles. Residual EB was removed via butanol extraction. An equal volume of water-saturated n-butanol was added to the reaction mixture, followed by brief vortexing and centrifugation at 450 g for 5 s. The upper organic phase was discarded. This step was repeated 2–3 times. The aqueous phase was then transferred to a fresh tube, mixed with two volumes of cold absolute ethanol and 1/10 volume of 3 M sodium acetate (pH 5.2), and incubated at –20 °C for at least 30 min. DNA was pelleted by centrifugation at 12,000 g for 10 min, washed twice with 70% ethanol, air-dried, and resuspended in nuclease-free water. The circularized DNA products were analyzed by 5% TBC-PAGE (acrylamide/bis = 29:1) in tris-borate-calcium buffer (90 mM Tris base, 90 mM boric acid, 10 mM CaCl_2_, pH 8.2) at 120 V for 120 min.

### Topoisomerase treatments of plasmid and minicircle DNA

For the Topoisomerase I reactions, 500 ng of negatively supercoiled plasmid DNA was incubated with 1 µL (5 U) Topoisomerase I (Novoprotein) at 37°C for 30 min, followed by heat inactivation at 65°C for 20 min. For relaxation of minicircle DNA, 100 ng of underwound 340-bp minicircle DNA prepared using 12 µg/mL ethidium bromide was treated under the same conditions. For DNA gyrase reactions, purified GyrA and GyrB proteins (each at 1 µM) were pre-assembled at 30°C for 30 min. Relaxed plasmid DNA (300 ng) or relaxed minicircle DNA prepared without ethidium bromide (150 ng) was incubated with increasing amounts of the assembled DNA gyrase complex (1, 3, 6, or 10 µL) at 37°C for 1 h in a reaction buffer containing 35 mM Tris-HCl (pH 7.5), 24 mM KCl, 4 mM MgCl_2_, 2.5 mM DTT, 5 mM spermidine, 0.1% BSA, 5% glycerol, and 1 mM ATP. Reactions were terminated by adding 1 µL proteinase K and incubating at 55 °C for 10 min. Plasmid DNA products were resolved on 0.8% TAE agarose gels and visualized by ethidium bromide staining. Minicircle DNA products were resolved by 5% TBC–PAGE and visualized by ethidium bromide staining.

### Reconstitution of the AtCas9-sgRNA-DNA ternary complex

The dAtCas9–sgRNA–DNA ternary complex was assembled and purified in two steps. First, dAtCas9 and sgRNA were incubated to form ribonucleoprotein (RNP) complexes, which were purified by size-exclusion chromatography (SEC). Purified RNPs were incubated with either annealed B-form DNA substrates or underwound minicircle DNA, followed by another round of SEC to isolate the fully assembled ternary complexes.

For the B-form DNA complex, dAtCas9, sgRNA and dsDNA were mixed at a molar ratio of 1:3:3. dAtCas9 was first incubated with annealed sgRNA at a molar ratio of 1:3 in Buffer 16A (20 mM HEPES, pH 7.5, 50 mM NaCl, 20 mM MgCl_2_, 0.5 mM DTT) at 50 °C for 40 min. The RNP mixture was then purified using a HiLoad 16/600 Superdex 200 pg column (Cytiva) and eluted in Buffer 16B (20 mM HEPES, pH 7.5, 150 mM NaCl, 20 mM MgCl_2_). Peak fractions corresponding to the RNP complex were pooled and concentrated to ∼250 µL using a centrifugal filter with a 50 kDa molecular weight cutoff (Millipore). Annealed dsDNA (TS: NTS = 1:3 molar ratio) was then incubated with purified RNP complexes in Buffer 16A at 50 °C for 40 min. The reaction was subjected to a second round of SEC on the same column equilibrated with Buffer 16B. Fractions corresponding to the ternary complex peak were pooled and concentrated to ∼1.0 mg/mL, as determined by NanoDrop, for subsequent cryo-EM sample preparation.

The dAtCas9-sgRNA-underwound minicircle DNA ternary complex was assembled using the same two-step procedure. To ensure complete saturation of all binding sites in underwound DNA, an excess of RNP was used (underwound DNA: dAtCas9: sgRNA = 1:41:123). Complex incubation and SEC purification were carried out under the same conditions as described above. Fractions corresponding to the ternary complex peak were pooled and concentrated to ∼0.4 mg/mL, as determined by NanoDrop, for subsequent cryo-EM sample preparation.

### Cryo-EM sample preparation and data collection

For the dAtCas9-sgRNA-B-form DNA complex, a total volume of 3.5 μL of the purified sample was applied onto glow-discharged (40 s under H_2_/O_2_ condition) EM grids (Quantifoil R1.2/1.3 Au300). The samples were left to stand for 60 s under 100% humidity at 8°C, and then blotted for 3 s with the bolt force of 3, using a Vitrobot Mark IV (FEI). For the dAtCas9-sgRNA-underwound minicircle DNA complex, 3.5 μL of the purified sample was applied onto glow-discharged EM grids (Quantifoil R1.2/1.3 Cu300, 2nm C), then immediately blotted for 3 s (blot force 3) under the same environmental conditions. Blotted grids were immediately plunge-frozen in liquid ethane. All grids were stored in liquid nitrogen until data collection.

For the dAtCas9-sgRNA-B-form DNA complex, cryo-EM datasets were collected on a Titan Krios electron microscope (Thermo Fisher Scientific) equipped with a K3 Summit direct electron detector (Gatan) at 300 kV accelerating voltage, using SerialEM software^31^ (version 3.7). The pixel size was 0.82 Å at a nominal magnification of ×29,000 and the defocus ranged from −1.0 to −2.5 μm. Each movie consisted of 40 frames with the exposure time of 2 s, and the total dose was 60 e^-^ Å^-2^ with a dose rate of 20 e^-^ pixel^-1^ s^-1^ (Supplementary Table 1).

For the dAtCas9-sgRNA-underwound DNA complex (CATA PAM, TATA PAM and TTGA PAM), cryo-EM datasets were collected on a Titan Krios G4 electron microscope (Thermo Fisher Scientific) equipped with a Gatan K3 direct electron detector and a Quantum energy filter (Gatan), operated at 300 kV accelerating voltage and using EPU software (Thermo Fisher Scientific). The pixel size was 0.84 Å at a nominal magnification of ×105,000 and the defocus ranged from -1.4 to -1.8 μm. Each movie consisted of 40 frames acquired over a 2 s exposure, with a total dose of 50 e^-^ Å^-2^ (Supplementary Table 1).

### Cryo-EM data processing

The cryo-EM data were processed using CryoSPARC^32^ (version 4.7.1). All the data sets for the four complexes were processed under the same pipeline as following, respectively. Firstly, the raw movies were imported and subjected to beam-induced motion correction using patch motion correction^33^ and fitted locally-variable CTF landscapes using patch CTF estimation^34^. Junk movies were removed manually. And then blob picker and template picker were used for particle picking. The initial particles were extracted and bad particles were removed through multiple rounds of 2D classification. The picked particles were subjected to several rounds of *ab initio* reconstruction to generate 4-6 models. The particles for the best-performing model were selected and subjected to heterogeneous refinement, with the best-performing model and three junk models as references. Then the output good particles were subjected to several rounds of heterogeneous refinement using the same strategy. The final target particles were further applied for homogenous refinement, non-uniform refinement and local refinement, generating density maps with indicated global resolutions at a gold-standard Fourier shell correlation (GSFSC) of 0.143.

For the dAtCas9-sgRNA-B-form DNA complex, 10,187 movies were collected, and 302,040 particles were finally obtained for generating the density map with a resolution of 3.15 Å (Extended data Fig. 2c-e and Supplementary Table 1). For the dAtCas9-sgRNA-underwound DNA complex (CATA PAM), 5,487 untilted and 2,378 30°-tilted movies were collected, and 452,994 particles were finally obtained for generating the density map with a resolution of 2.41 Å (Extended data Fig. 5f-h and Supplementary Table 1). For the dAtCas9-sgRNA-underwound DNA complex (TATA PAM), 6,718 untilted and 2,090 30°-tilted movies were collected, and 457,333 particles were finally obtained for generating the density map with a resolution of 2.65 Å (Extended data Fig. 6a-c and Supplementary Table 1). For the dAtCas9-sgRNA-underwound DNA complex (TTGA PAM), 7,028 untilted and 1,449 30°-tilted movies were collected, and 1,023,398 particles were finally obtained for generating the density map with a resolution of 2.89 Å (Extended data Fig. 6d-f and Supplementary Table 1).

### Structural model building, refinement and analysis

The structure of Nme1Cas9-sgRNA-DNA complex (PDB: 6KC7)^35^ was used as an initial model for fitting into the EM density maps using UCSF ChimeraX^36^ (version 1.9). The model was subjected to manual adjustment and rebuilding in Coot^37^ (version 0.9.8.95) to fix alternate residues and nucleotides. Then the coordinates of dAtCas9-sgRNA-B-form DNA complex and dAtCas9-sgRNA-underwound DNA complexes (CATA PAM, TATA PAM, and TTGA PAM) were refined for several rounds by Coot and real-space refinement with secondary structure constrains in Phenix^38^ (version 1.20.1). The model statistics were validated via MolProbity^39^. Final refinement statistics were provided in Extended data Table 1. UCSF ChimeraX^36^ (version 1.9) and PyMOL (Version 3.1, Schrödinger, LLC.) were used for structural figure preparation in this paper.

### Calculation of DNA bending angle

To quantify the bending angle of the target-strand DNA between the PAM duplex and the DNA:sgRNA heteroduplex, the global helical axes of the full-length PAM duplex and the distal 17-bp region of the DNA:sgRNA heteroduplex were defined using principal component analysis (PCA). PCA was performed on the step-position coordinates (Px, Py, Pz) derived from x3DNA analysis^40^. The inter-helical bending angle was then calculated as the arccosine of the dot product between the two normalized helical axis vectors.

## Data availability

The atomic coordinates for dAtCas9-sgRNA-B-form DNA complex, dAtCas9-sgRNA-underwound DNA complex (CATA PAM), dAtCas9-sgRNA-underwound DNA complex (TATA PAM), and AtCas9-sgRNA-underwound DNA complex (TTGA PAM) have been deposited in the Protein Data Bank (PDB) with the accession codes 9WAD, 21DZ, 21EA and 9WAC, respectively. The cryo-EM density maps for dAtCas9-sgRNA-B-form DNA complex, dAtCas9-sgRNA-underwound DNA complex (CATA PAM), dAtCas9-sgRNA-underwound DNA complex (TATA PAM), and AtCas9-sgRNA-underwound DNA complex (TTGA PAM) have been deposited in the Electron Microscopy Data Bank (EMDB) with the accession codes EMD-65812, EMD-67605, EMD-67606 and EMD-65811, respectively.

---

**Extended Data Figure 1.**
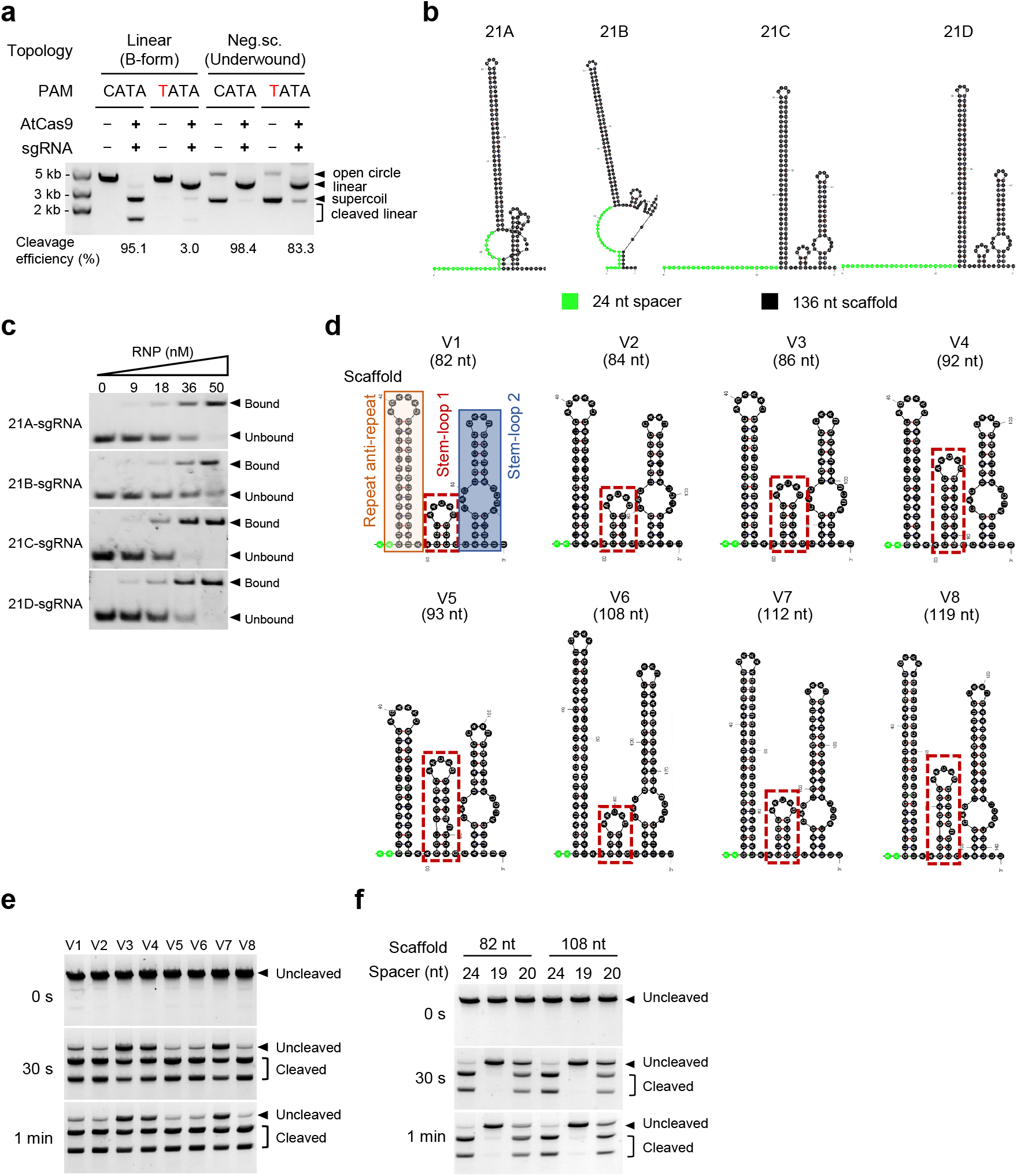
Optimization of sgRNA structure. **a**, *In vitro* cleavage assay of AtCas9 on WT PAM and mutant PAM under different DNA topologies. **b**, Schematic structure of full-length sgRNAs with different spacer sequences. The 24-nt spacer is highlighted in green, and the 136-nt scaffold is shown in black. **c**, EMSA of various sgRNA-dAtCas9 RNPs with corresponding substrate DNAs. **d**, Schematic of different sgRNA scaffold structures. The repeat:anti-repeat region was truncated by 14-nt, from 48-nt to 34-nt. The stem region of stem-loop 2 was truncated by 12-nt. Stem-loop 1, highlighted with a red box, was engineered with an extended stem and alternative secondary structures. **e**, *In vitro* cleavage of sgRNAs with modified scaffold. Cleavage performed at room temperature for the indicated times. **f**, *In vitro* cleavage of sgRNAs with variable spacer length.

**Extended Data Figure 2.**
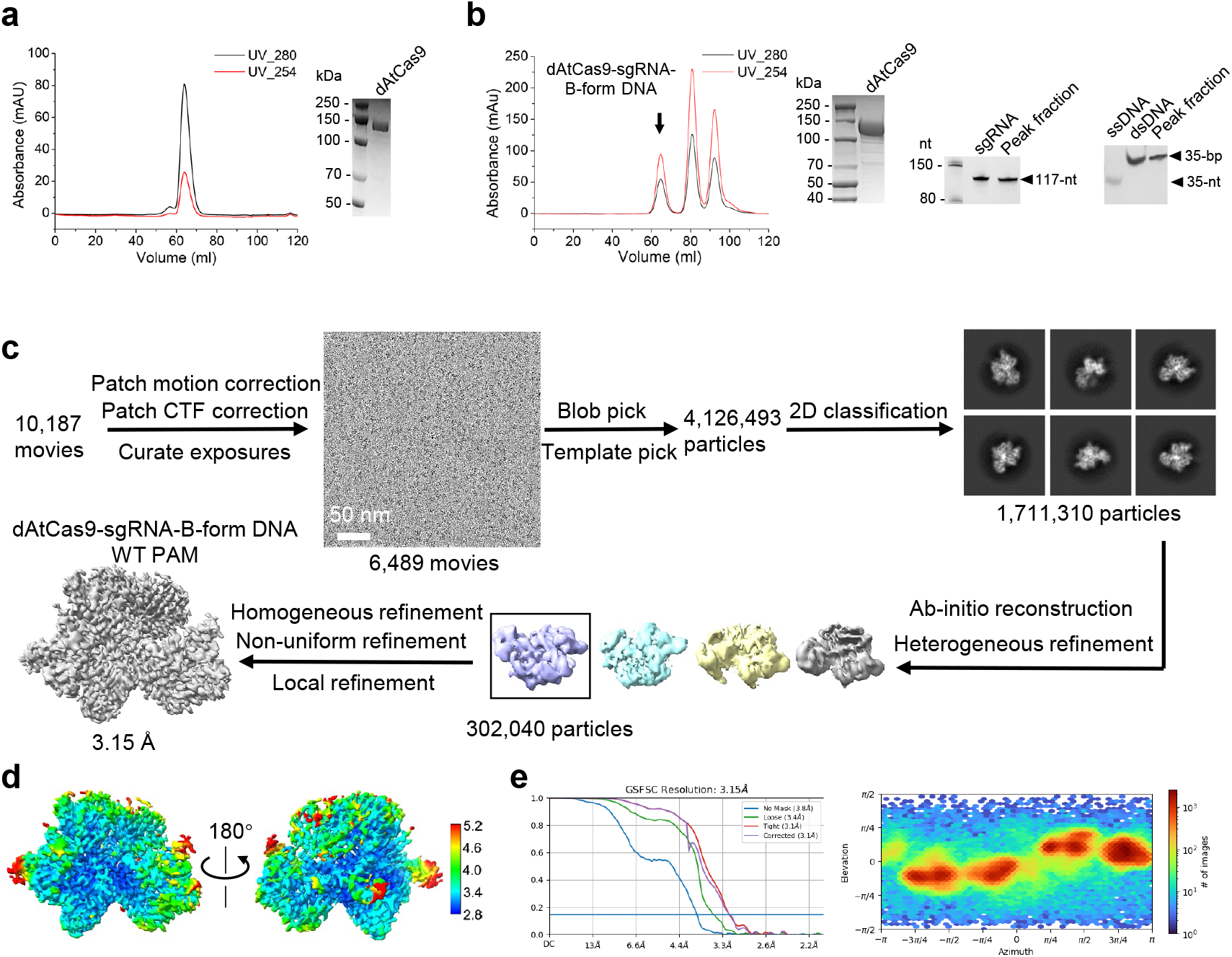
Cryo-EM sample preparation and data processing workflow. **a**, Size-exclusion chromatography of the dAtCas9 and SDS-PAGE analysis. **b**, Size-exclusion chromatography of the dAtCas9-sgRNA-B-form DNA complex, followed by SDS-PAGE, Urea-PAGE, and TBE-PAGE analyses of AtCas9, sgRNA, and dsDNA, respectively. The black arrow indicates the peak corresponding to the ternary complex. **c**, Cryo-EM data processing pipeline for dAtCas9-sgRNA-B-form DNA complex. **d**, Resolution ranges of the 3D density map for dAtCas9-sgRNA-B-form DNA complex. **e**, Fourier shell correlation (FSC) curves of the 3D density maps and the euler angle distributions of particles for dAtCas9-sgRNA-B-form DNA complex.

**Extended Data Figure 3.**
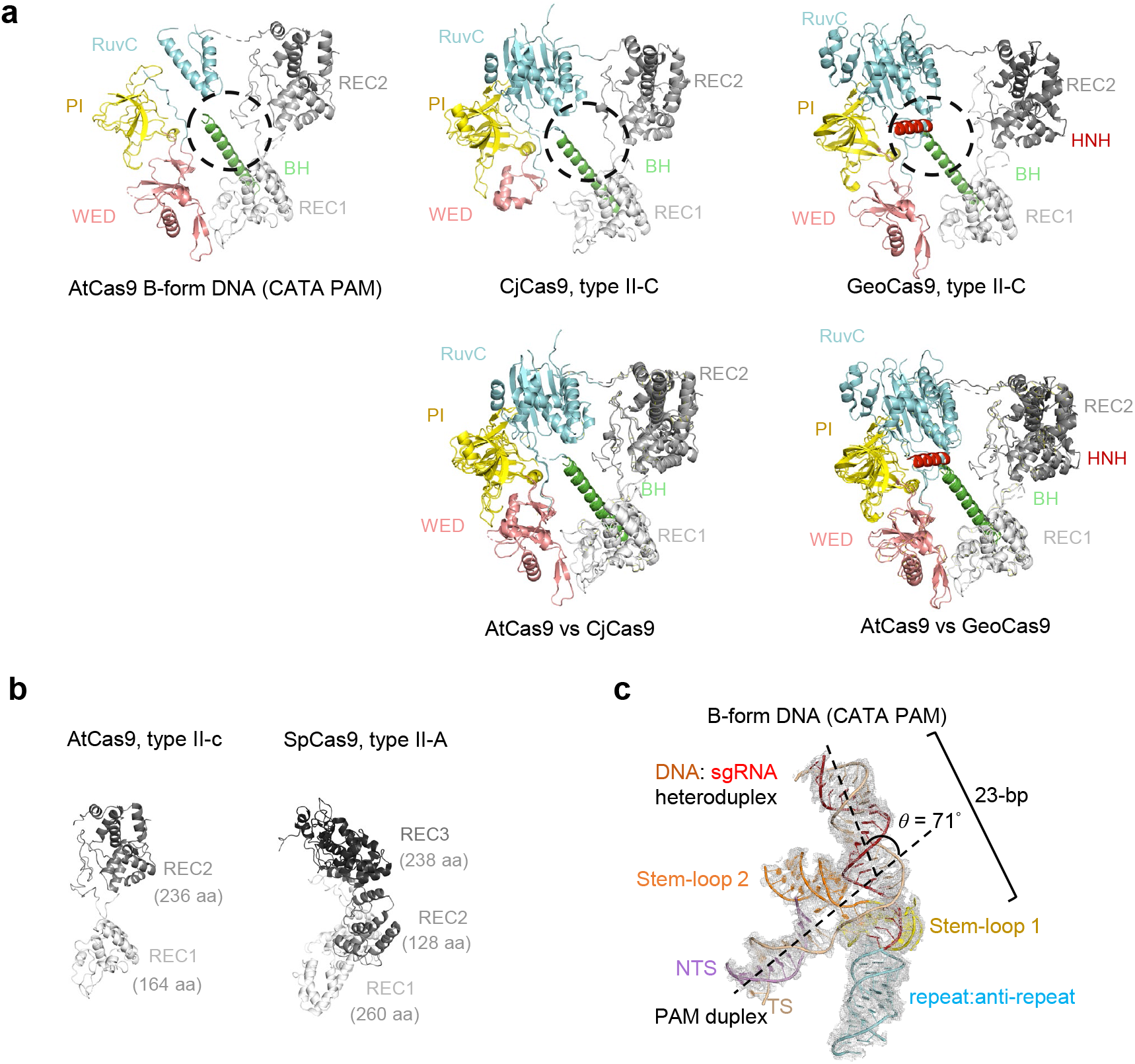
Structural comparison of the Cas9 orthologs. **a**, Structural comparison of AtCas9 upon binding to B-form DNA (PDB: 9WAD) with CjCas9 (PDB: 5X2G) and GeoCas9 (PDB: 8UZA). **b**, REC domain comparison of AtCas9 (PDB: 9WAD) and SpCas9 (PDB: 5F9R). **c**, Structure of the DNA-sgRNA in dAtCas9-sgRNA-B-form DNA complex. *θ* denotes the bending angle between the PAM-proximal and PAM-distal duplexes.

**Extended Data Figure 4.**
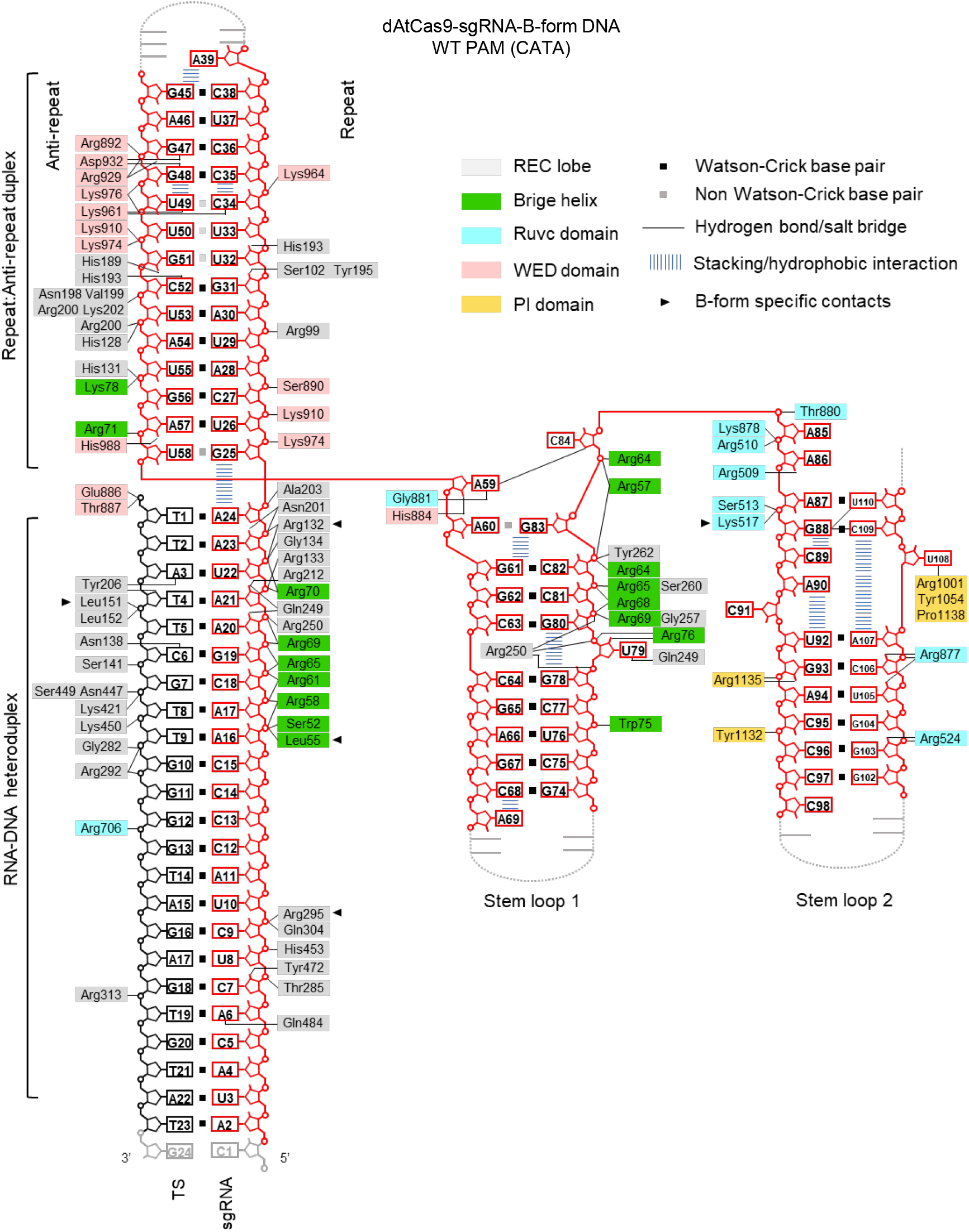
Schematic representation of sgRNA-target DNA recognition by AtCas9 in the B-form DNA complex. Residues interacting with the sgRNA and its target DNA are colored according to their respective domains. B-form-specific interacting residues are indicated with black triangles. Disordered regions are shown as gray dashed lines.

**Extended Data Figure 5.**
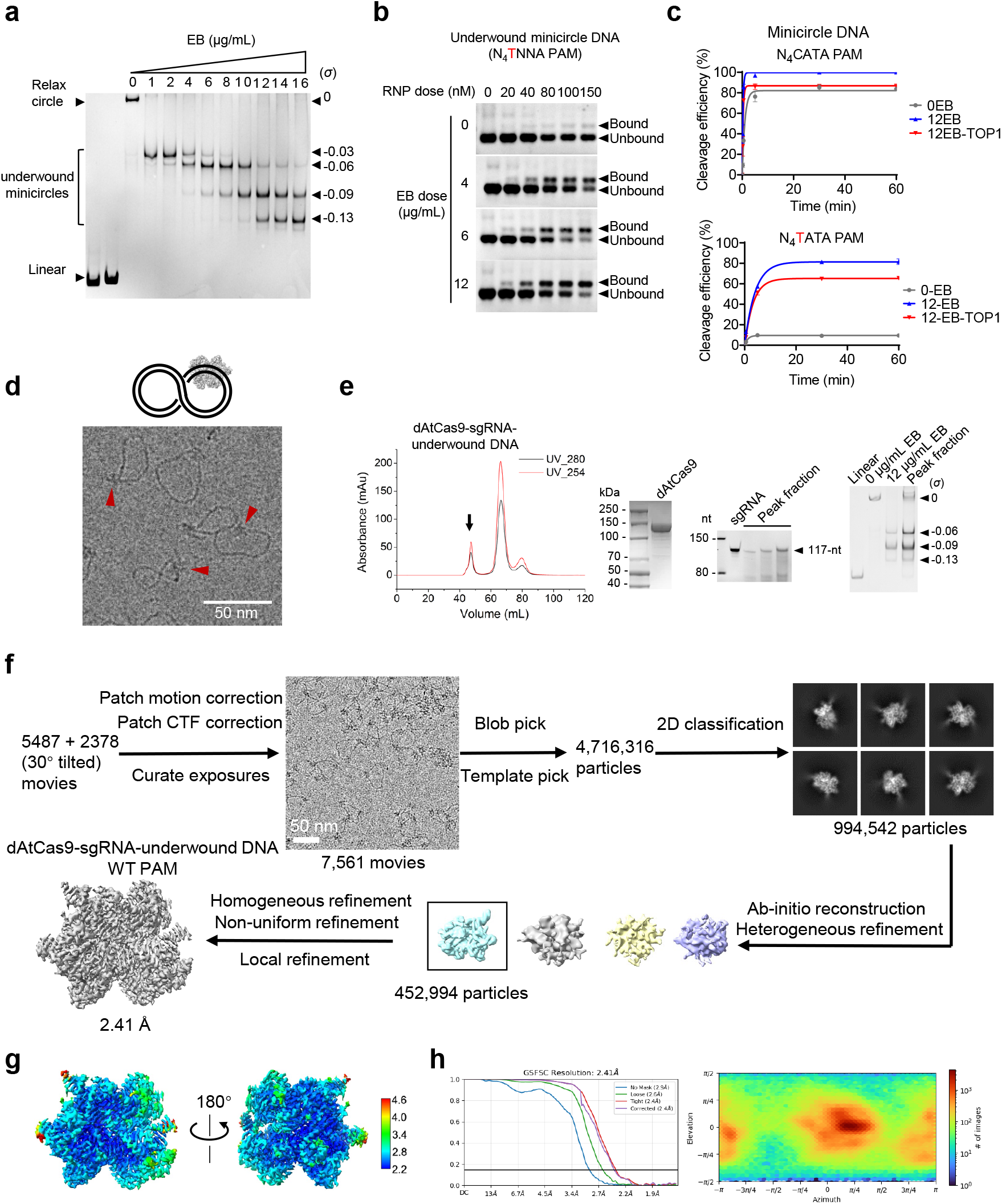
Characterization of 340-bp negatively supercoiled minicircle DNA, and their Cryo-EM sample preparation and data processing. **a**, Gel electrophoresis analysis of EB-induced topoisomers of 340-bp circular DNA. σ denotes the superhelical density. **b**, EMSA analysis of dAtCas9-RNP bound to the indicated 340-bp circular DNA topoisomers generated with varying EB dosages. **c**, *In vitro* cleavage kinetics of TOP1-relaxed minicircles. Underwound minicircles generated by EB treatment (12EB) were relaxed using TOP1 prior to cleavage assays. Substrates bearing WT PAM CATA or MUT1 PAM TATA were tested. TOP1, topoisomerase I. Data are represented as mean ± SD and fitted with non-linear regression. **d**, Schematic and representative cryo-EM micrographs of dAtCas9-RNP bound to underwound minicircle DNA with a single target site. RNP particles bound to minicircle DNA are indicated by red arrows. **e**, Size-exclusion chromatography of the dAtCas9-sgRNA-underwound DNA (WT PAM) complex, followed by SDS-PAGE, Urea-PAGE, and TBC-PAGE analyses of AtCas9, sgRNA, and minicircle DNA, respectively. The black arrow indicates the peak corresponding to the ternary complex. **f**, Cryo-EM data processing pipeline for dAtCas9-sgRNA-underwound DNA (WT PAM) complex. **g**, Resolution ranges of the 3D density map for dAtCas9-sgRNA-underwound DNA (WT PAM) complex. **h**, FSC curves of the 3D density maps and the euler angle distributions of particles for dAtCas9-sgRNA-underwound DNA (WT PAM) complex.

**Extended Data Figure 6.**
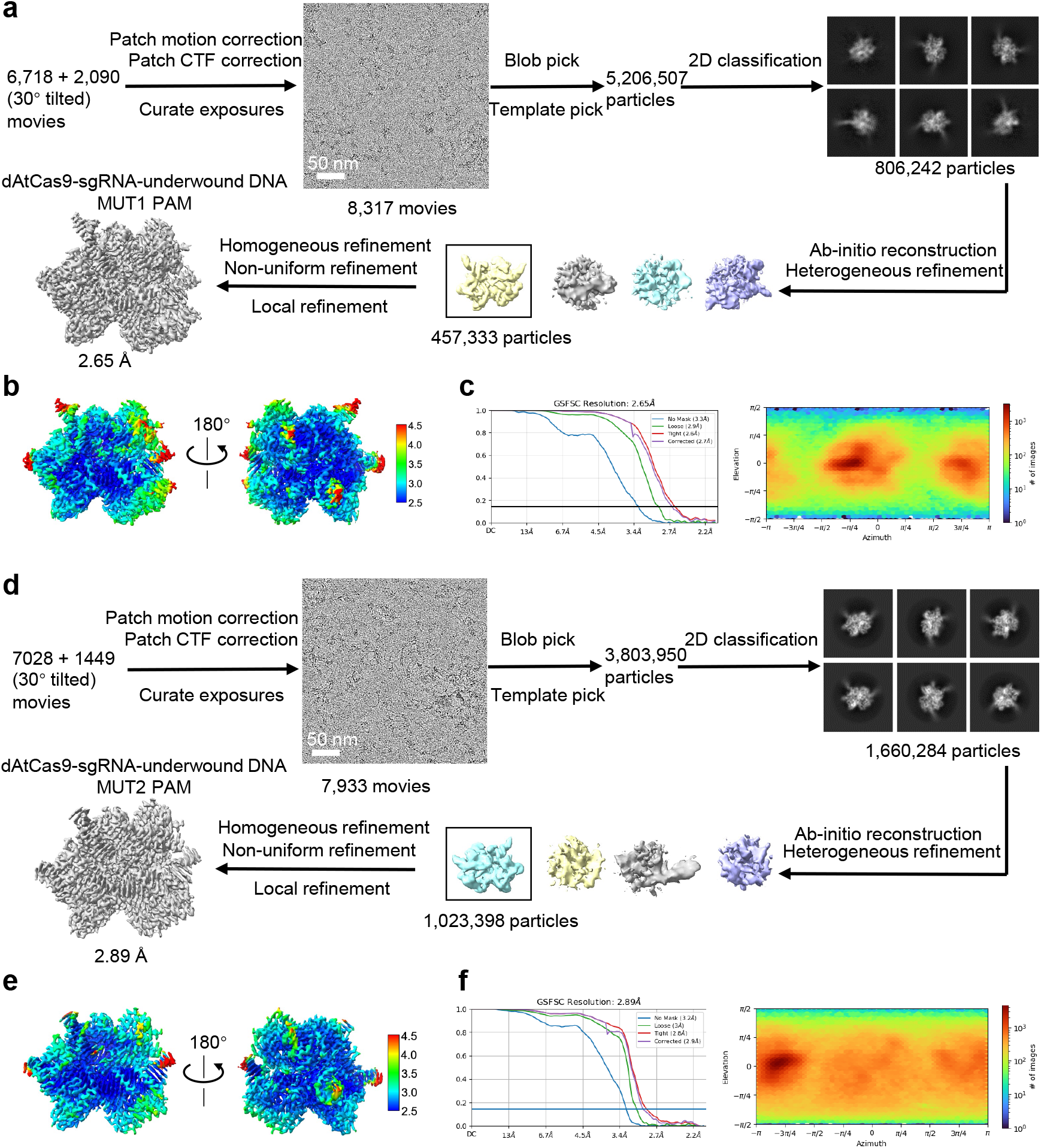
Cryo-EM data processing workflow for the underwound DNA complex containing the MUTANT PAMs. **a**, Cryo-EM data processing pipeline for dAtCas9-sgRNA-underwound DNA (MUT1 PAM) complex. **b**, Resolution ranges of the 3D density map for dAtCas9-sgRNA-underwound DNA (MUT1 PAM) complex. **c**, FSC curves of the 3D density maps and the euler angle distributions of particles for dAtCas9-sgRNA-underwound DNA (MUT1 PAM) complex. **d**, Cryo-EM data processing pipeline for dAtCas9-sgRNA-underwound DNA (MUT2 PAM) complex. **e**, Resolution ranges of the 3D density map for dAtCas9-sgRNA-underwound DNA (MUT2 PAM) complex. **f**, FSC curves of the 3D density maps and the euler angle distributions of particles for dAtCas9-sgRNA-underwound DNA (MUT2 PAM) complex.

**Extended Data Figure 7.**
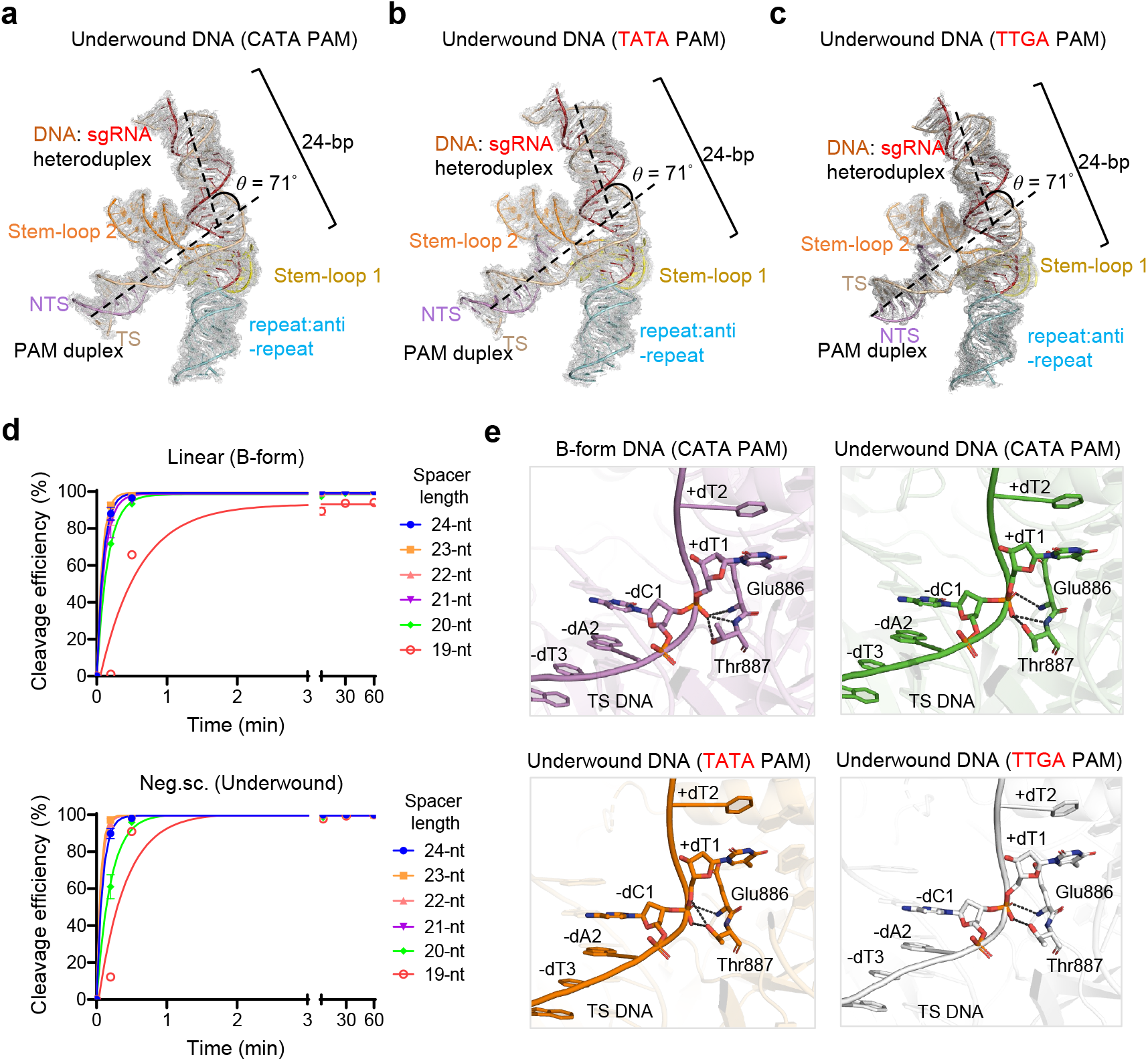
Characterization of DNA-sgRNA hybrids in underwound DNA complex containing the WT PAM and MUTANT PAMs. **a-c**, Structure of the DNA-sgRNA heteroduplex in dAtCas9-sgRNA-underwound DNA complex containing WT PAM (**a**), MUT1 PAM (**b**) and MUT2 PAM (**c**). *θ* denotes the bending angle between the PAM-proximal and PAM-distal duplexes. **d**, Kinetic cleavage assay of sgRNAs with variable spacer lengths. Data are represented as mean ± SD and fitted with non-linear regression. **e**, Close-up views of the phosphate lock loop in the B-form and three underwound DNA complexes.

**Extended Data Figure 8.**
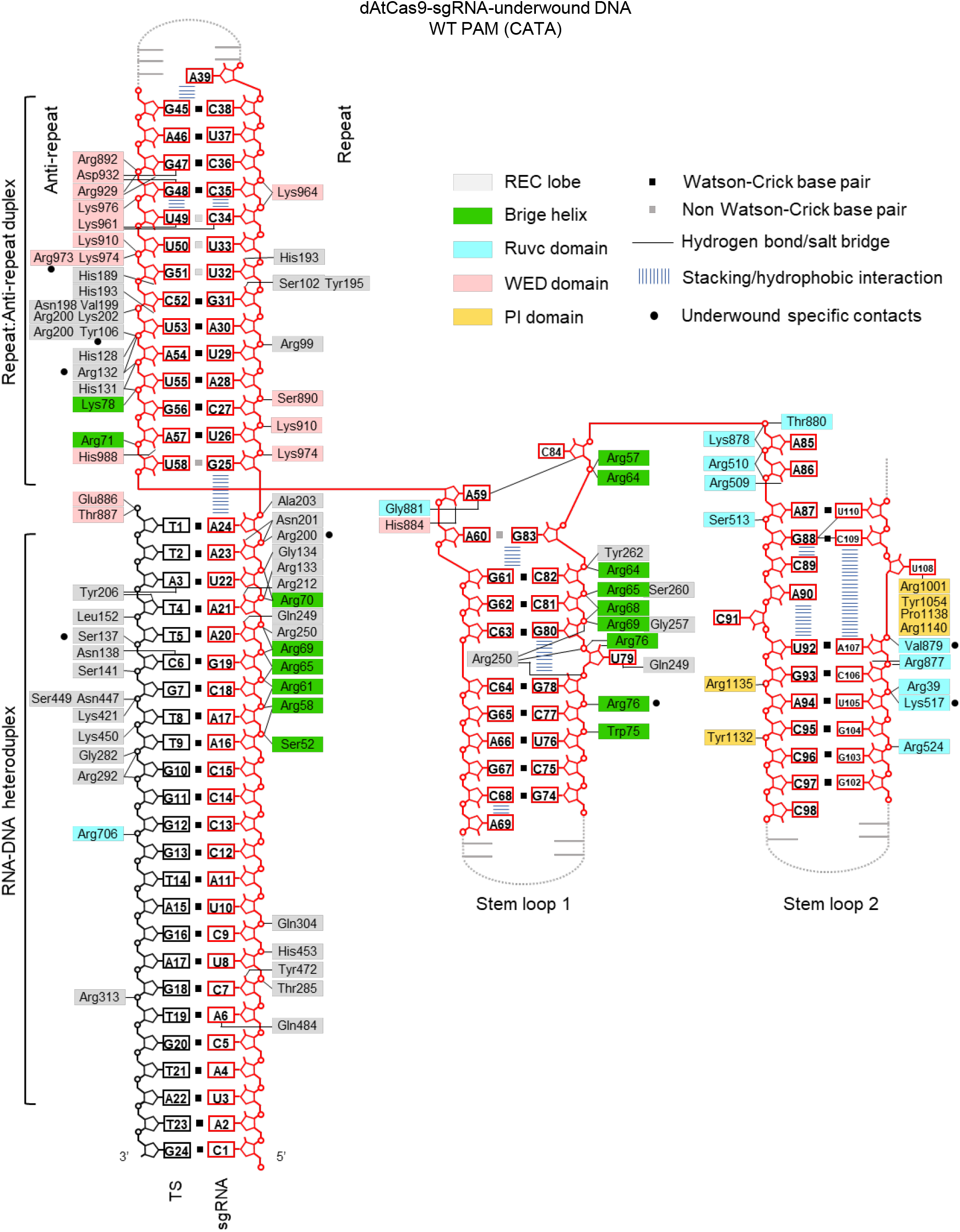
Schematic representation of sgRNA-target DNA recognition by AtCas9 in the underwound DNA complex containing the WT PAM. Residues interacting with the sgRNA and its target DNA are colored according to their respective domains. Underwound-specific interacting residues are indicated with black circles. Disordered regions are shown as gray dashed lines.

**Extended Data Figure 9.**
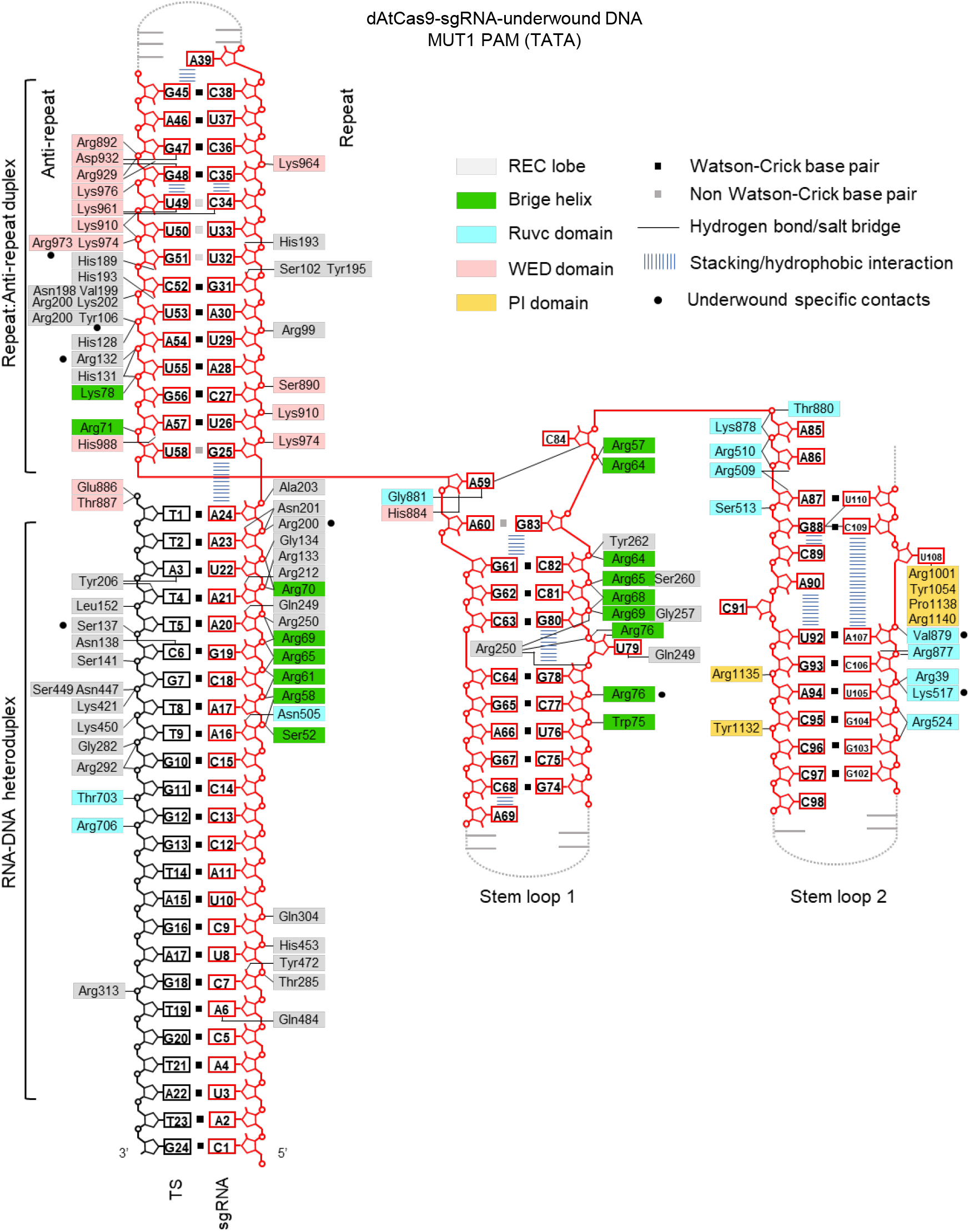
Schematic representation of sgRNA-target DNA recognition by AtCas9 in the underwound DNA complex containing the MUT1 PAM. Residues interacting with the sgRNA and its target DNA are colored according to their respective domains. Disordered regions are shown as gray dashed lines.

**Extended Data Figure 10.**
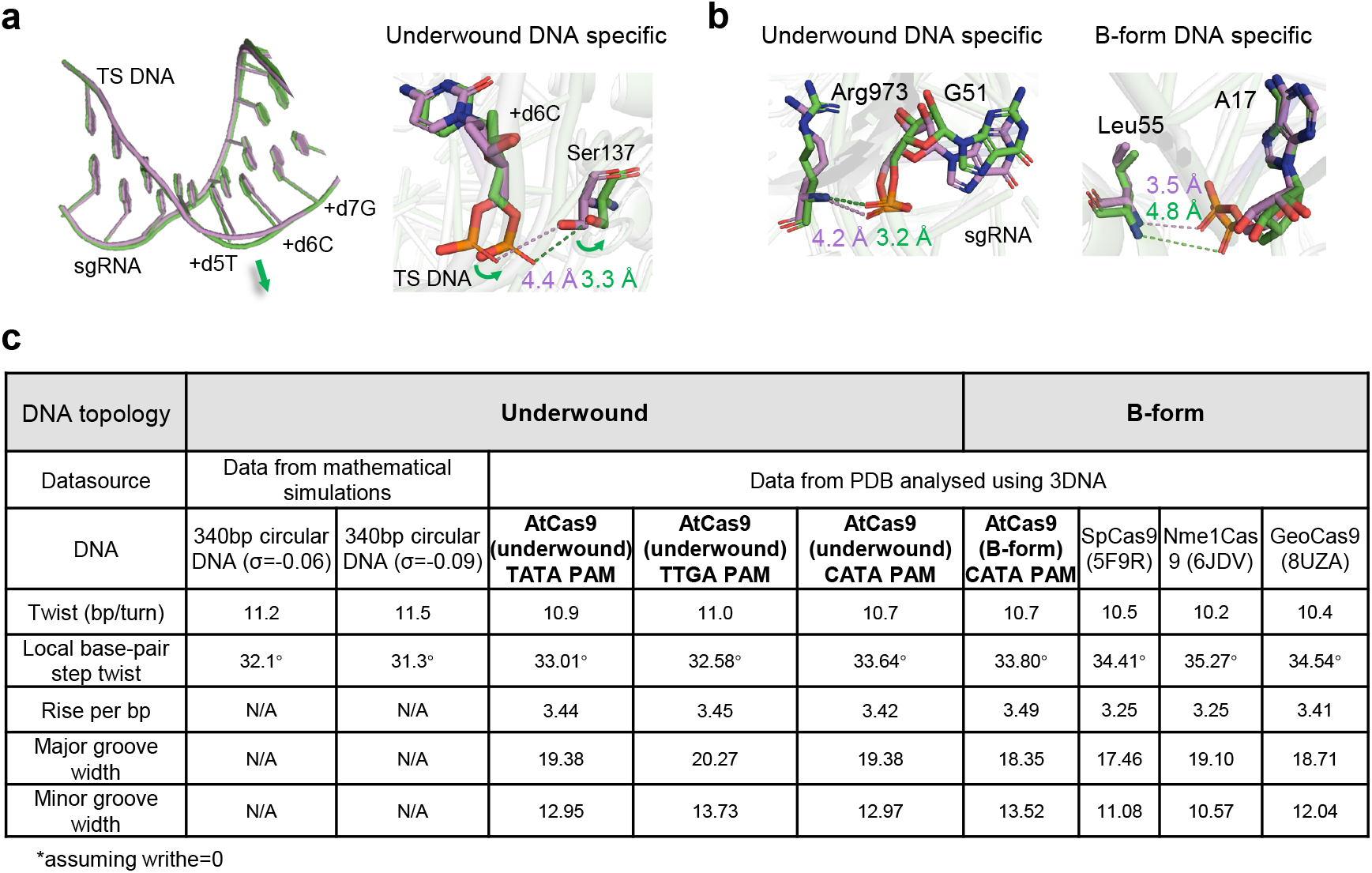
Comparison of topology-specific interaction sites and DNA topological parameters. **a**, Structural comparison of the R-loops in B-form DNA (violet) and underwound DNA (green) complexes containing the WT PAM. The expansion direction of the underwound DNA relative to B-form DNA is indicated by green arrow. Close-up view of the superposition of TS DNA-specific interacting residues are shown on the right. **b**, Close-up views of the superposition of representative sgRNA-specific interacting residues in the underwound and B-form DNA complexes containing the WT PAM. **c**, 340-bp minicircle models with superhelical densities of σ = -0.06 and -0.09 were generated assuming that torsional stress is entirely accommodated by twist in the absence of writhe (Wr = 0). DNA structural parameters of underwound and B-form AtCas9 complexes, together with SpCas9 (PDB 5F9R), Nme1Cas9 (PDB 6JDV) and GeoCas9 (PDB 8UZA), were analyzed using 3DNA^40^.

**Extended Data Figure 11.**
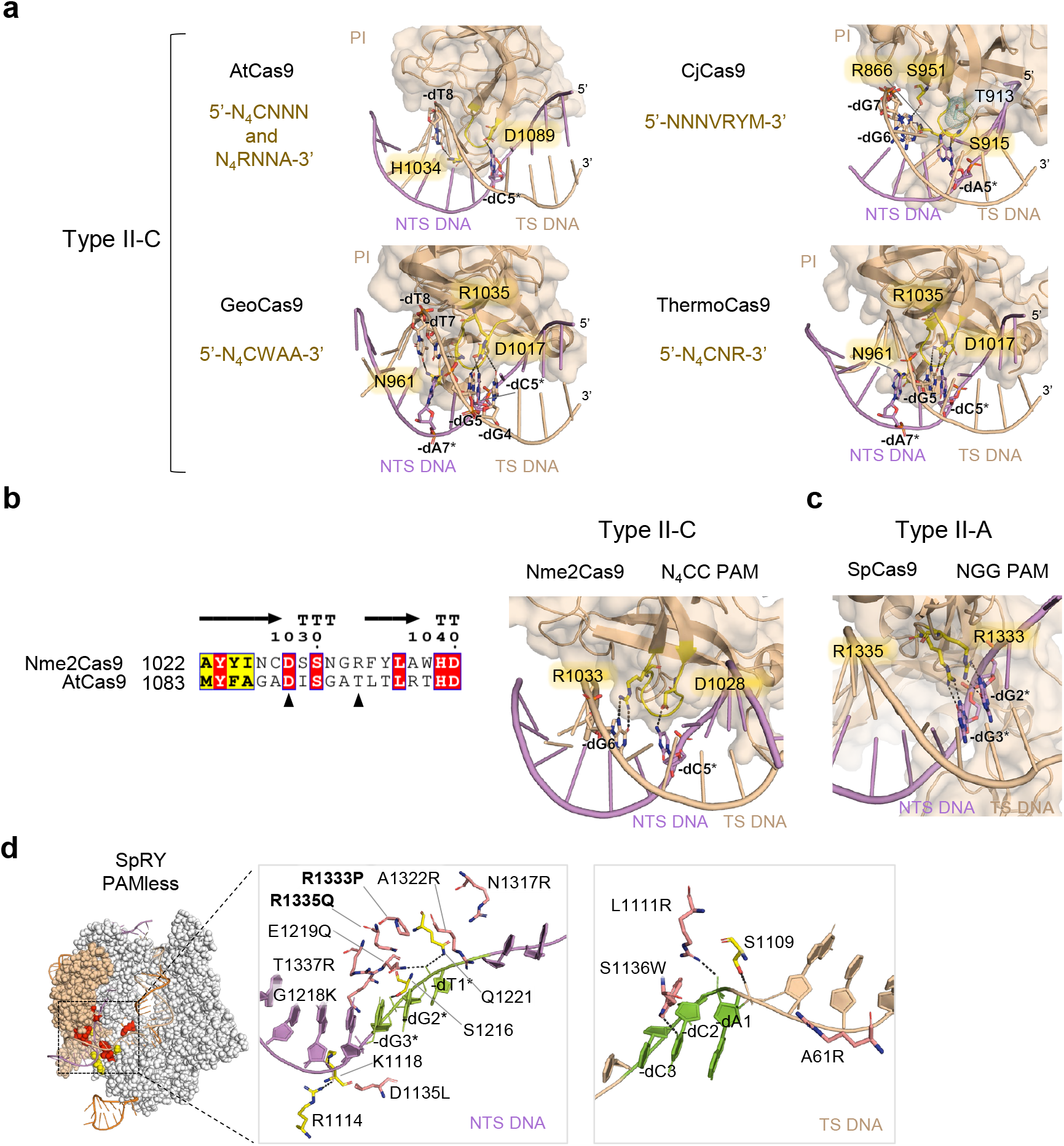
PAM engagement and base-specific recognition among Cas9 Orthologs. **a**, Overall structures and base-specific interaction residues of the PI domain and PAM duplex in Cas9 orthologs, including AtCas9 (PDB: 9WAD), GeoCas9 (PDB: 8UZA), CjCas9 (PDB: 5X2G) and ThermoCas9 (PDB: 9AR6). PI domains are represented by surface and cartoon, with interacting residues highlighted in yellow. In CjCas9, Thr913, which enforces preference at the fourth PAM position via steric clash with the methyl group of dT4*, is shown as cyan dots. NTS, non-target strand; TS, target strand. **b**, Sequence alignment of the PAM-interacting regions of AtCas9 and Nme2Cas9 (left), with an enlarged view of PAM recognition in Nme2Cas9 (right) (PDB: 6JE3). **c**, Enlarged view of PAM recognition in SpCas9 (PDB: 5F9R). **d**, Overview of the SpRY (PDB: 8SPQ) with a close-up view of the PAM-interacting region, highlighting mutations relative to SpCas9. Backbone-interacting residues for the TS and NTS strands within the PAM duplex (yellow) and nearby mutated residues (red) are indicated.

**Supplementary Figure 1.**
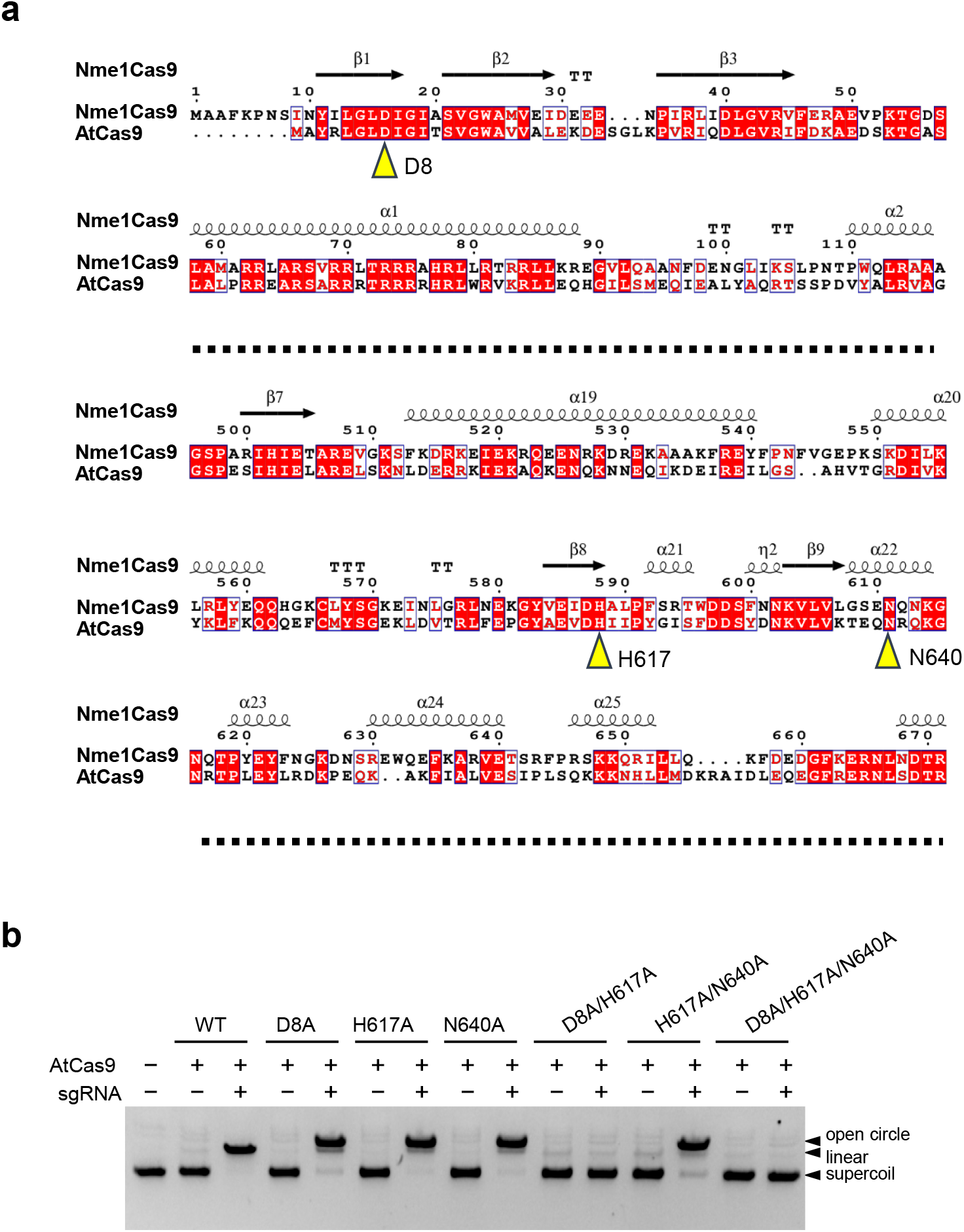
Screening of catalytic sites in AtCas9. **a**, Sequence alignment of AtCas9 and Nme1Cas9. Catalytic residues in the HNH and RuvC domains of Nme1Cas9 were aligned to identify corresponding key residues in AtCas9 (highlighted in yellow). **b**, *In vitro* cleavage of AtCas9 variants carrying mutations in the predicted catalytic residues.

**Supplementary Figure 2.**
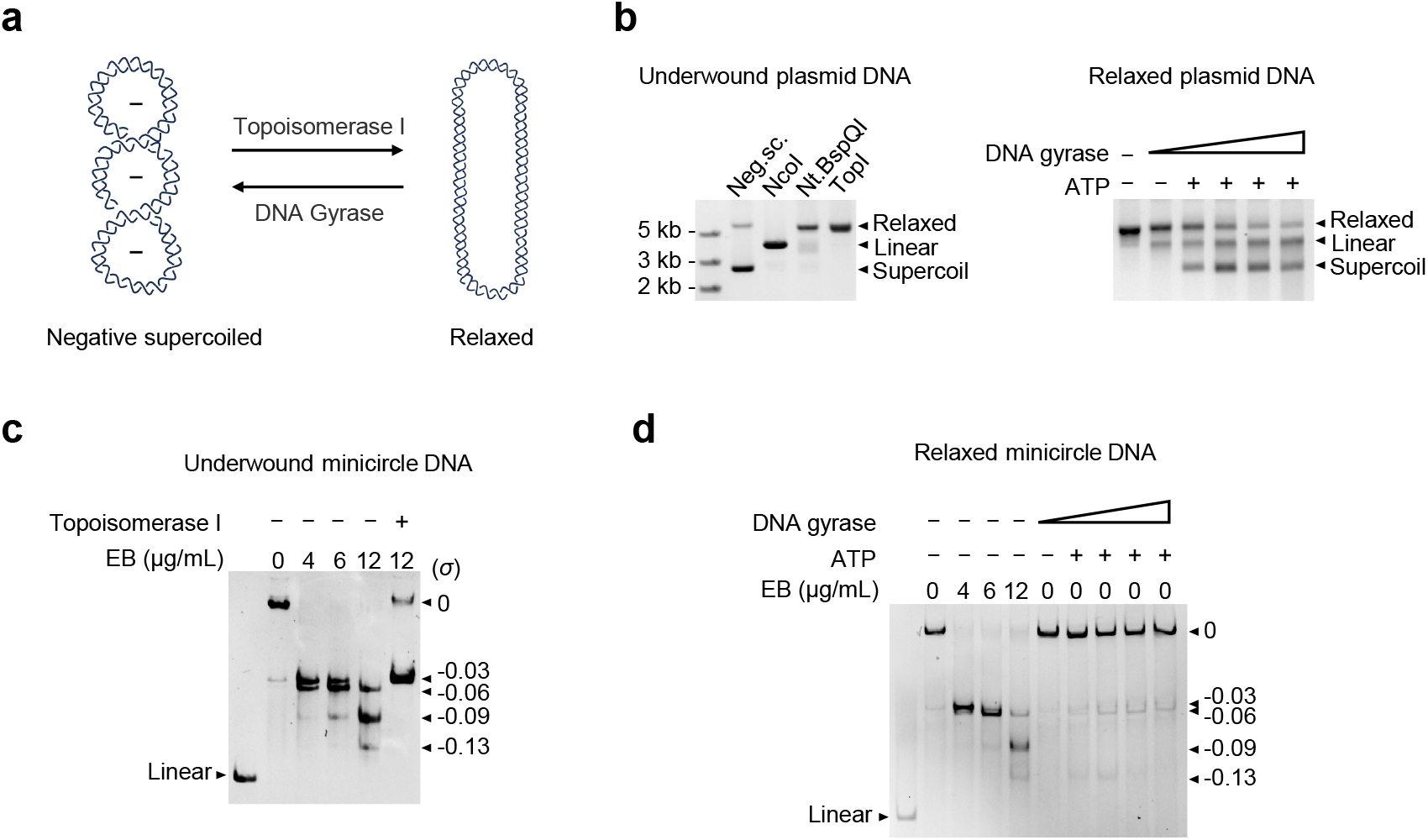
Characterization of the topological states of minicircle DNA using DNA topoisomerases. **a**, Illustration of DNA topological changes induced by Topoisomerase I or DNA gyrase. **b**, Gel electrophoresis analysis of plasmid DNA topological states following treatment with topoisomerase I (TOP I) or DNA gyrase. Linear: plasmids were linearized using NcoI. Relaxed: plasmids were treated with TOP I. Neg. Sc.: negatively supercoiled plasmids. For TOP I reaction, negative supercoiled plasmids served as substrates; for DNA gyrase reaction, relaxed plasmids served as substrates. **c**, Gel electrophoresis analysis of 340-bp underwound minicircle DNA prepared with 12 µg/mL EB after treatment with TOP I. **d**, Gel electrophoresis analysis of underwound 340-bp minicircle DNA prepared using DNA gyrase.

**Supplementary Figure 3.**
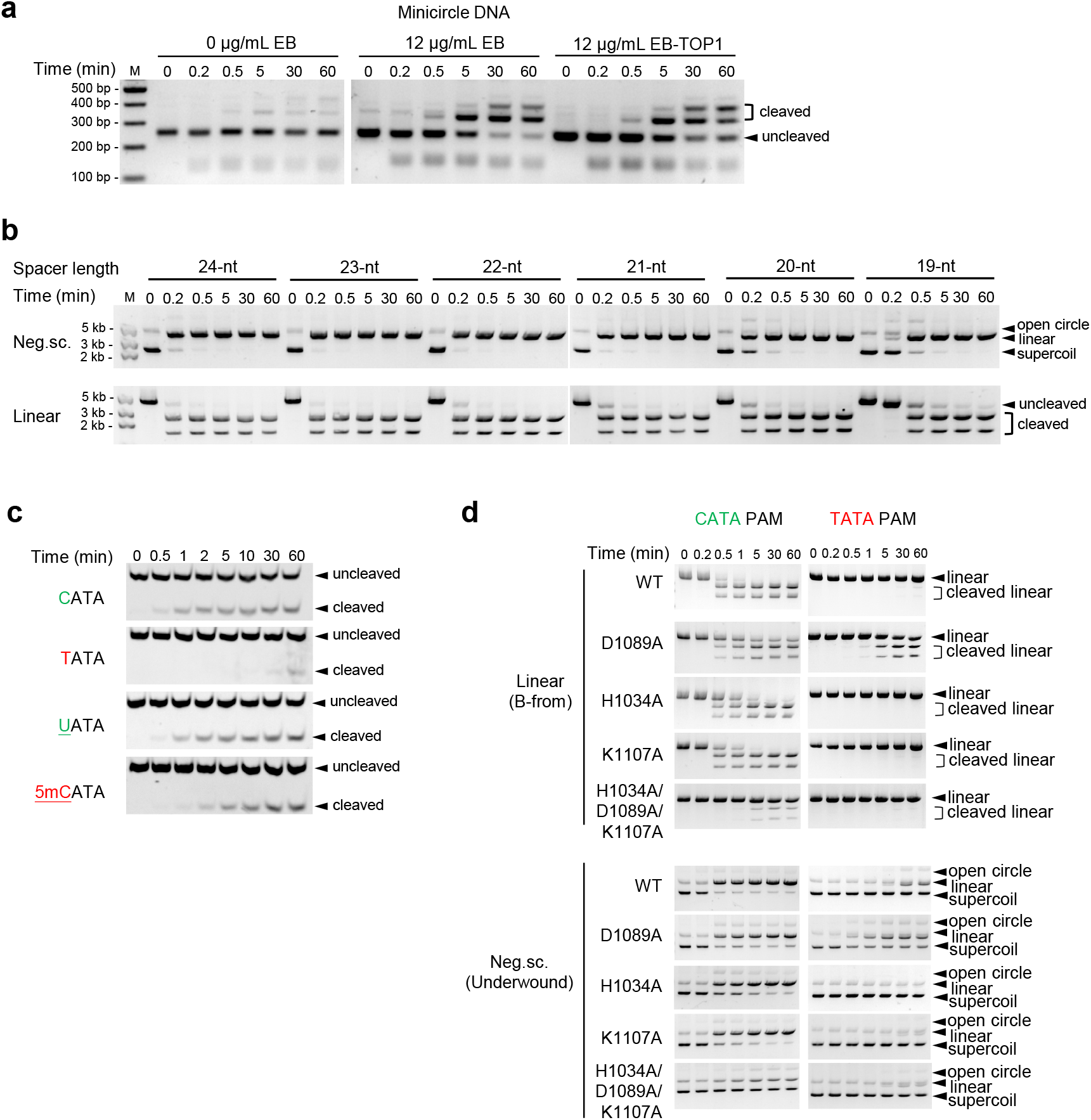
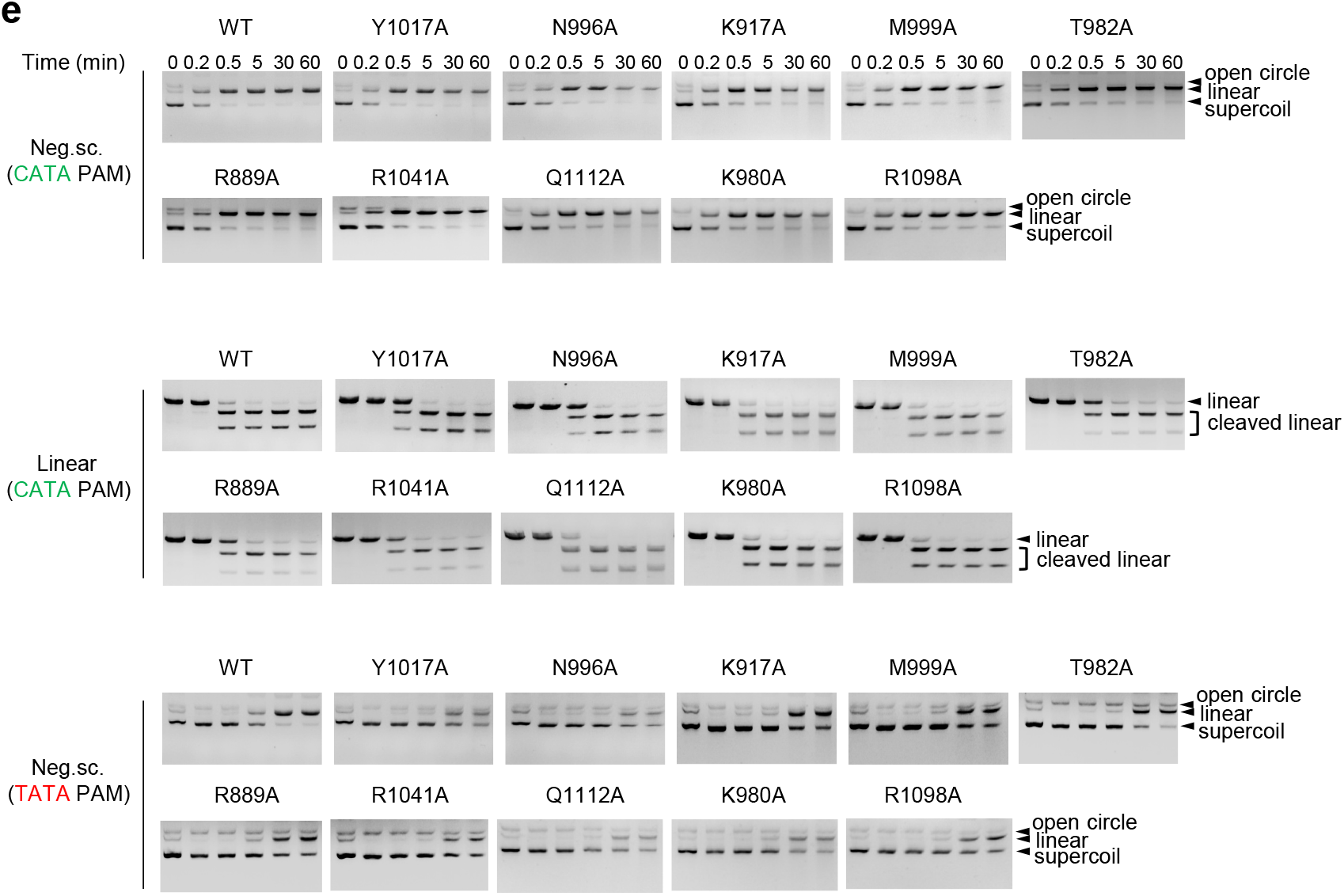
Kinetic analysis of DNA cleavage by AtCas9. **a**, *In vitro* cleavage kinetics of TOP1-relaxed minicircles (raw data for Extended Data Fig. 5c). RNPs were incubated with different topoisomers of minicircle DNA for the indicated times at 55 °C. **b**, *In vitro* cleavage kinetics assay of sgRNAs with variable spacer lengths (raw data for Extended Data Fig. 7d). RNPs (100 nM) were incubated with negatively supercoiled plasmids or linearized plasmids (2 nM) for the indicated times at 55 °C. **c**, *In vitro* cleavage kinetics assays of dAtCas9 with different PAM variants: CTAA, TATA, UATA, and 5mCTAA (raw data for Fig. 4b). 40 nM RNPs and 20 nM 5’ FAM-labeled oligonucleotides (50 bp) were incubated for indicated time at 55°C. **d**, *In vitro* cleavage kinetics assays of AtCas9 bearing indicated mutations on WT PAM (CATA) and mutant PAM (TATA) under different DNA topologies (raw data for Fig. 4f). RNPs (20 nM) were incubated with negatively supercoiled plasmids or linearized plasmids (2 nM) for the indicated times at 55 °C. **e**, *In vitro* cleavage kinetics assays of AtCas9 bearing indicated mutations on WT PAM (CATA) and mutant PAM (TATA) under different DNA topologies (raw data for Fig. 5e). RNPs (100 nM) were incubated with negatively supercoiled plasmids or linearized plasmids (2 nM) for the indicated times at 55 °C.

**Supplementary Table 1.** Cryo-EM data collection and processing, model refinement and validation statistics.

|  | #1 dAtCas9-sgRNA-B-form DNA<br>(CATA PAM)<br>(EMDB-65812)<br>(PDB 9WAD) | #2 dAtCas9-sgRNA-underwound DNA<br>(CATA PAM)<br>(EMDB-67605)<br>(PDB 21DZ) | #3 dAtCas9-sgRNA-underwound DNA<br>(TATA PAM)<br>(EMDB-67606)<br>(PDB 21EA) | #4 dAtCas9-sgRNA-underwound DNA<br>(TTGA PAM)<br>(EMDB-65811)<br>(PDB 9WAC) |
| --- | --- | --- | --- | --- |
| Data collection and processing |  |  |  |  |
| Magnification | 29,000 | 105,000 | 105,000 | 105,000 |
| Voltage (kV) | 300 | 300 | 300 | 300 |
| Electron exposure (e<sup>-</sup>/Å<sup>2</sup>) | 60 | 50 | 50 | 50 |
| Defocus range (µm) | -1.0~-2.5 | -1.4~-1.8 | -1.4~-1.8 | -1.4~-1.8 |
| Pixel size (Å) | 0.82 | 0.84 | 0.84 | 0.84 |
| Symmetry imposed | C1 | C1 | C1 | C1 |
| Initial particle images (No.) | 4,126,493 | 4,716,316 | 5,206,507 | 3,803,950 |
| Final particle images (No.) | 302,040 | 452,994 | 457,333 | 1,023,398 |
| Map resolution (Å) | 3.15 | 2.41 | 2.65 | 2.89 |
| FSC threshold | 0.143 | 0.143 | 0.143 | 0.143 |
| Map resolution range (Å) | 2.81-3.83 | 2.15-2.68 | 2.31-2.92 | 2.54-3.53 |
| Refinement |  |  |  |  |
| Initial model used (PDB code) | 6KC7 | 6KC7 | 6KC7 | 6KC7 |
| Model-Map CC (mask) | 0.73 | 0.86 | 0.81 | 0.75 |
| Map sharpening *B* factor (Å<sup>2</sup>) | 106.7 | 77.5 | 83.3 | 106.9 |
| Model composition |  |  |  |  |
| Non-hydrogen atoms | 8590 | 9747 | 9750 | 9502 |
| Protein residues | 701 | 842 | 839 | 827 |
| Nucleotide residues | 141 | 144 | 145 | 138 |
| *B* factors (Å<sup>2</sup>) |  |  |  |  |
| Protein | 50.07 | 40.90 | 61.40 | 42.76 |
| Nucleotide | 74.11 | 22.33 | 26.94 | 21.57 |
| R.m.s. deviations |  |  |  |  |
| Bond lengths (Å) | 0.003 | 0.004 | 0.004 | 0.004 |
| Bond angles (°) | 0.557 | 0.624 | 0.605 | 0.618 |
| Validation |  |  |  |  |
| MolProbity score | 1.96 | 1.47 | 1.60 | 1.86 |
| Clashscore | 10.56 | 4.45 | 5.60 | 8.25 |
| Poor rotamers (%) | 0.00 | 0.14 | 0.42 | 0.56 |
| Ramachandran plot (%) |  |  |  |  |
| Favored (%) | 93.67 | 96.36 | 95.72 | 93.66 |
| Allowed (%) | 6.33 | 3.64 | 4.28 | 6.34 |
| Disallowed (%) | 0.00 | 0.00 | 0.00 | 0.00 |

